# Stable isotope–assisted computational mass spectrometry reveals root-specific alkaloids in *Glycyrrhiza* species

**DOI:** 10.64898/2026.05.05.722977

**Authors:** Keita Sawai, Yoshimasa Todoroki, Shohei Nakamukai, Yuki Matsuzawa, Keiichi Noguchi, Toshiyo Kato, Tetsuya Mori, Amit Rai, Masami Yokota Hirai, Hiroshi Tsugawa

**Affiliations:** Department of Biotechnology and Life Science, Tokyo University of Agriculture and Technology, 2-24-16 Naka-cho, Koganei, Tokyo 184-8588, Japan; RIKEN Center for Sustainable Resource Science, 1-7-22 Suehiro-cho, Tsurumi-ku, Yokohama, Kanagawa 230-0045, Japan; Graduate School of Pharmaceutical Sciences, Kyoto University, 46-29 Yoshida-Shimoadachi-cho, Sakyo-ku, Kyoto 606-8501, Japan; Scientific-materials Creating Open Plaza, Tokyo University of Agriculture and Technology, 2-24-16 Naka-cho, Koganei, Tokyo 184-8588, Japan; Department of Crop Sciences, University of Illinois Urbana-Champaign, 1201 Gregory Dr, Urbana IL 61801, USA

**Author notes:** **Corresponding Author** Hiroshi Tsugawa.

## Abstract

Licorice (*Glycyrrhiza*) is a medicinal plant widely used in approximately 70% of traditional Japanese Kampo formulations and is known to produce a wide array of specialized metabolites with diverse pharmacological properties. Although hundreds of metabolites have been reported, the overall chemical diversity of *Glycyrrhiza* remains poorly characterized. Here, using mass spectrometry data obtained from fully ^13^C-labeled leaves and roots of *Glycyrrhiza uralensis* and *Glycyrrhiza glabra*, we determined the carbon number, followed by molecular formula and substructure prediction in combination with MS/MS similarity-based molecular networking. After excluding redundant ions, including isotopic peaks, adducts, and in-source fragments, we extracted 3,060 unique metabolite features with assigned carbon numbers. Among these, substructure information was assigned to 1,015 features (33%) across the four plant tissues, revealing the tissue-specific metabolome profiles. Furthermore, we discovered five previously unreported alkaloids, homopipecolic acid-conjugated flavonoids, in the roots of *G. uralensis* and *G. glabra*, and *Glycine max*, another member of the Fabaceae family. Two of these structures were validated using nuclear magnetic resonance spectroscopy. We further proposed a biosynthetic route involving a spontaneous reaction between 1-piperideine and malonyl glycoside substrates and confirmed the formation of the conjugated product using authentic standards.

## Introduction

Licorice (*Glycyrrhiza spp*.), a globally important medicinal and industrial plant used in pharmaceuticals, foods, and other consumer products, is one of the most frequently utilized crude drugs in Japan and is included in more than 70% of Kampo formulations [1]. Although licorice is often discussed in terms of its representative constituents, such as glycyrrhizin and glycyrrhizic acid, licorice-derived products contain diverse classes of specialized metabolites, including triterpenoid saponins and flavonoids, whose combined composition is thought to be important for their biological properties [1,2]. Despite extensive phytochemical research, a comprehensive system-level understanding of licorice metabolite diversity is lacking. Database-scale summaries and literature-linked resources contain hundreds of *Glycyrrhiza*-associated metabolites [3]. However, these reports are fragmented across tissues, studies, and analytical scopes, complicating efforts to connect chemistry with biosynthetic pathways and comparative quality traits [4,5].

Plants frequently localize specialized metabolites to specific organs, tissues, and cell types [6,7,8]. This phenomenon is indicative of tissue-specific biosynthesis, transport, and storage, which are integral components of environmental and biotic adaptation [6,7,8]. For licorice, this metabolite distribution is of particular relevance, given that “licorice” as a crude drug is predominantly derived from roots and rhizomes, whereas aerial parts are underutilized despite evidence that they contain distinct constituents and bioactivities, such as antimicrobial and antioxidant-related activities reported for leaf/aerial extracts [9,10]. Thus, tissue-resolved metabolomics, which compares aerial and subterranean organs, is imperative not only for cataloging compounds but also for elucidating the pathways and metabolite families that are preferentially expressed in specific tissues. This approach enables a biologically interpretable framework of specialized licorice metabolism [4]. Furthermore, licorice is not unique. *Glycyrrhiza uralensis* and *Glycyrrhiza glabra* are the principal plant species used worldwide. Multiomics and metabolomics studies have indicated substantial chemical differences among *Glycyrrhiza* species, suggesting that species identity is a core determinant of metabolite profiles and pathway regulation [4,5,11]. Therefore, a comparative design analyzing both *G. uralensis* and *G. glabra* under aligned conditions is essential to illuminate conserved and species-specific metabolomes and to establish a reference framework for downstream gene-metabolite analyses and cross-species interpretation [4,5].

Liquid chromatography–tandem mass spectrometry (LC–MS/MS) is a widely utilized technique in plant metabolomics owing to its sensitivity, mass accuracy, and comprehensive fragmentation information for elucidating metabolite structures [12]. However, structural annotation remains a significant impediment; untargeted LC–MS/MS studies frequently assign confident identities to only a small percentage of detected peaks, prompting approaches that incorporate orthogonal constraints to enhance structural inference for unknowns [13]. Stable isotope labeling is a particularly powerful constraint because the carbon-shift observation between labeled and unlabeled plants indicates that the carbon number can narrow the candidate formulae and structures for unknown metabolites and increase confidence in computational annotation workflows [14,15,16]. In our previous study [14], we introduced a stable isotope-assisted cheminformatics workflow implemented in MS-DIAL to enable carbon number determination and structural characterization across diverse plant species and tissues. Importantly, this evaluation included licorice tissues from *G. uralensis* and *G. glabra* [14]. Concurrently, owing to the heightened sensitivity of plant metabolomes to cultivation and microenvironmental variations, the employment of paired labeled/unlabeled designs necessitates meticulously matched growth conditions to circumvent the confounding of biological differences with labeling conditions [17,18]. In the present study, licorice plants were cultivated under closely matched conditions and comprehensively profiled to generate a tissue-resolved metabolome landscape focused on *Glycyrrhiza*, overcoming limitations of previous cross-species benchmarking analyses [14]. This dataset enabled systematic characterization of tissue-specific metabolic profiles and facilitated the discovery of previously unreported alkaloids.

## Materials and Methods

### Metabolite extraction

^13^C-labeled and unlabeled leaf, stem, and root samples of *G. uralensis, G. glabra*, and *Glycine max* were purchased from IsoLife (Wageningen, Netherlands). Metabolites were extracted from the labeled and unlabeled plant samples according to a previously described protocol [19]. For metabolite extraction, 50 μL of 80% methanol (MeOH) per 1 mg of dry sample, containing 2.5 μM lidocaine and 10-camphorsulfonic acid as internal standards, was added to suspend the powdered material. The suspension was homogenized using a mixer mill with zirconia beads for 7 min at 18 Hz and 4 °C. After centrifugation for 10 min, the supernatant was collected and filtered using an HLB μElution plate (Waters, Milford, MA, USA). The filtrate was transferred to an LC-MS vial fitted with an insert (Agilent Technologies, USA).

### LC-MS/MS condition

The experimental procedures were performed according to a previously described protocol [14]. The liquid chromatography (LC) system was an ExonLC system (SCIEX, Framingham, MA, USA). Metabolites were separated on an ACQUITY UPLC BEH C18 Column (100 × 2.1 mm:1.7 μm) (Waters, U.S.A). The column temperature was maintained at 40 °C. The mobile phases were (A) H_2_O with 0.1% formic acid and (B) acetonitrile with 0.1% of formic acid. A sample volume of 1 μL was injected. The separation was performed using the following gradient program: 0 min 0.5% (B), 0.1 min 0.5% (B), 10 min 80% (B), 10.1 min 99.5% (B), 12.0 min 99.5% (B), 12.1 min 0.5% (B), and 15.0 min 0.5% (B). The flow rate was set as follows: 0.3 mL/min at 0 min, 0.3 mL/min at 10 min, 0.4 mL/min at 10.1 min, 0.4 mL/min at 14.4 min, and 0.3 mL/min at 14.5 min. The temperature of the rack was maintained at 4 °C. MS detection was performed using a QTOF-MS (ZenoTOF 7600; SCIEX, Framingham, MA, USA). The parameters for positive and negative ion mode analyses were as follows: MS1 and MS2 mass ranges, *m/z* 50-1500; MS1 accumulation time, 200 ms; Q1 resolution, unit; MS2 accumulation time, 50 ms; maximum candidate ions, 10; ion source temperature, 450 °C; and CAD gas, 7. The following settings were used for positive/negative ion mode, independently: ion source gas 1, 50/50 psi; ion source gas 2, 50/50 psi; curtain gas, 35/35 psi; spray voltage, 5500/-4500 V; declustering potential, 80/-80 V; and collision energy, 35/-35 ± 15 eV.

### Data processing using MS-DIAL 4

Data processing was performed using MS-DIAL (version 4) [20]. Both MS1 and MS/MS data were processed as centroid data. For data collection, the MS1 mass tolerance was set to 0.01 Da and the MS2 mass tolerance to 0.025 Da. The peak detection parameters were set as follows: minimum peak height, 100 amplitude; mass slice width, 0.1 Da. An in-house spectral library constructed from public and commercially available databases, including MassBank and NIST, was used for metabolite annotation. The library consisted of 1,340,293 spectra corresponding to 41,922 compounds in the positive ion mode and 384,101 spectra corresponding to 26,493 compounds in the negative ion mode. Annotation was performed with the following parameters: retention time tolerance, 100 min; accurate mass tolerance (MS1), 0.01 Da; accurate mass tolerance (MS2), 0.05 Da; and identification score cutoff of 80%. For alignment, the retention time tolerance was set to 0.025 min, and the MS1 tolerance was set to 0.015 Da. All parameters that are not explicitly mentioned were maintained at their default settings.

### MS-FINDER parameter settings

The molecular formula was estimated using MS-FINDER [21], by incorporating the carbon number information (Fig. 1B). Default libraries were also used as data sources for the molecular formula prediction. The mass tolerance settings were set to ±5 ppm for MS1 and ±10 ppm for MS2. All other parameters were maintained at the default settings of MS-FINDER. Peaks with molecular formula prediction scores greater than three were successfully assigned.

**Figure 1.**
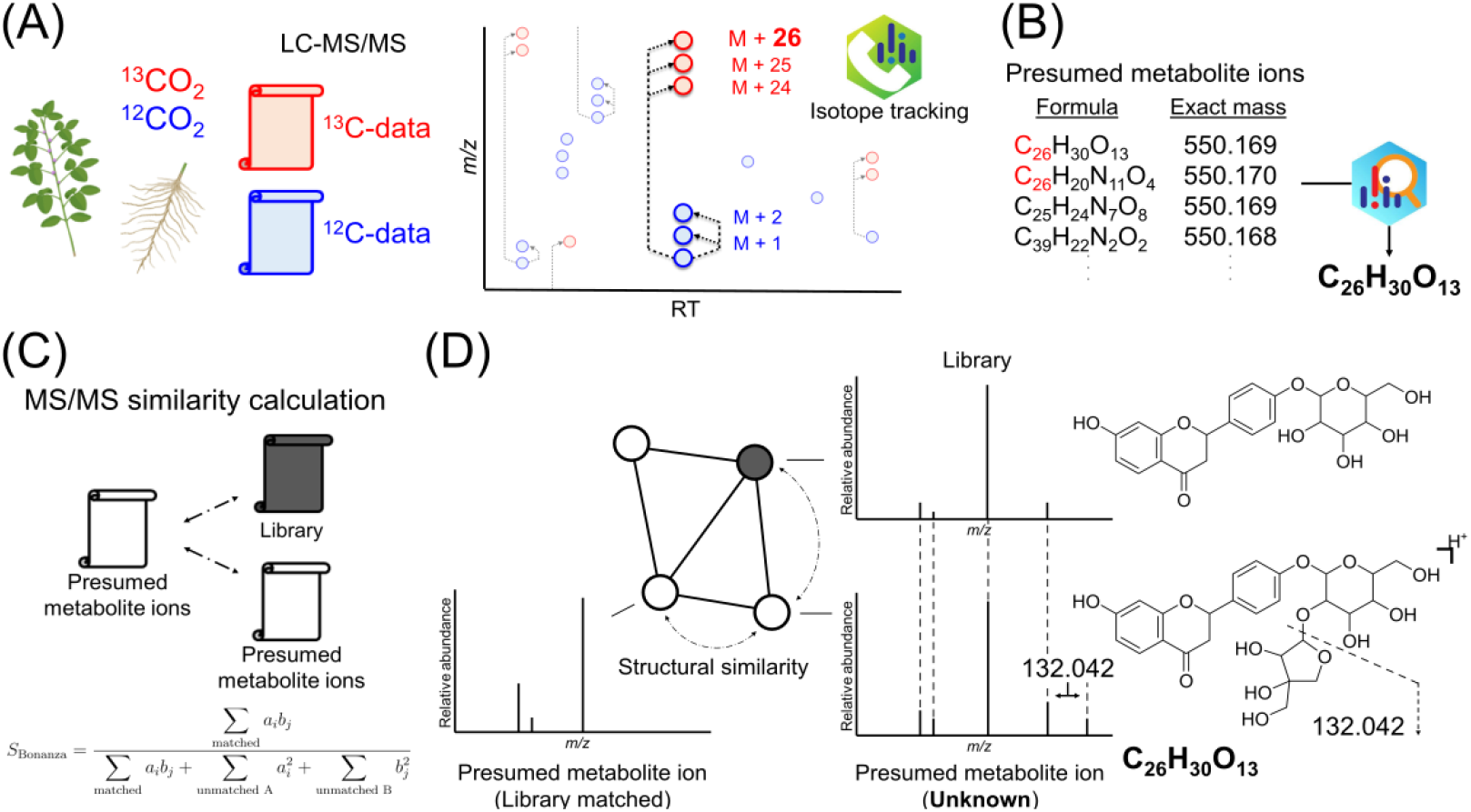
Overview of this study. (A) Estimation of carbon numbers and molecular formulas using stable isotope-labeled plants. LC-MS/MS data from labeled and unlabeled samples were integrated, and carbon numbers were estimated based on *m/z* shifts using the isotope-labeled tracking function on MS-DIAL 4. (B) Estimation of the molecular formula based on the carbon number information.

### Construction of MS/MS spectral similarity based molecular networks

Prior to network construction, the MS/MS spectra were preprocessed. Peaks within the *m/z* range of 0–10,000 were considered. Peaks with relative intensities below 0.1% of the base peak or absolute intensities below 50 were excluded. Fragment peaks with *m/z* values higher than those of the precursor ions were excluded. Product ion peaks were bound with a bin width of 1.0 Da. The squared root-transformed peak intensities were calculated and normalized such that the maximum intensity in the product-ion spectrum was set to 100. The isotopic peaks in the product ion spectra were not removed. Only peak pairs with precursor ion *m/z* differences of ≤400 Da were included in the analyses. The peaks were connected as edges in the network when they shared at least six matched fragment peaks and exhibited a Bonanza similarity score of ≥0.8. To limit network complexity, a maximum of 10 edges per node were retained and ranked according to their similarity scores.

### Isolation and purification of licorice alkaloids

Licorice roots (100 g; Tohoku, Uchida Wakanyaku, Ltd.) were ground into a fine powder using a high-speed mill (HS-15; Labonext Co., Ltd., Japan). The powdered sample was soaked in 1.9 L of 70% ethanol (EtOH) for 1 h at room temperature to obtain an extract. After filtration, the extract was concentrated under reduced pressure using a rotary evaporator (RE400; Yamato Scientific Co., Ltd., Japan). The dried extract was partitioned between water and CHCl_3_ and the aqueous phase was further extracted with *n*-BuOH. The *n*-BuOH fraction (6.85 g) was evaporated and separated on ODS flash column chromatography (Cosmosil® 75C18-OPN; Nacalai Tesque, Japan, φ 20 × 100 mm) with aqueous MeOH (stepwise 20%, 40%, 60%, 80%, and 100%) containing 0.1% acetic acid. The 40% MeOH fraction (1.47 g) was dissolved in 20 mL of MeOH and loaded into 2-mL aliquots onto a weak cation-exchange cartridge (Strata WCX, 5 g/20 mL; Phenomenex, USA) that had been preconditioned and equilibrated. After sequential washing with 60 mL of water and 60 mL of MeOH, the cartridge was eluted with 80 mL of MeOH containing 5% formic acid. The resulting eluates were combined to afford a final eluate of 92.23.

The eluate was evaporated to dryness, dissolved in 80% MeOH, and purified by reverse-phase HPLC using a Prominence system (Shimadzu, Japan) equipped with an autosampler (SIL-10AP), pump (LC-20AD), PDA detector (SPD-M20A), and a column oven (CTO-20A). The separation column was a Unison UK-C18 (250 mm × 10 mm i.d., 3 μm; Imtakt, Japan). The mobile phases were (A) H_2_O containing 0.1% formic acid and (B) acetonitrile containing 0.1% formic acid. Gradient elution was performed as follows: 0 min 0.5% (B), 0.1 min 0.5% (B), 5 min 20% (B), 25 min 20% (B), 26 min 99.5% (B), 31 min 99.5% (B), 32 min 0.5% (B), and 37 min 0.5% (B). The flow rate was 4 mL/min, and the column temperature was maintained at 40 °C. Two peaks detected by PDA at approximately 20 min were collected separately, affording mixtures of (iv) flavanone base; 2O: O-Hex; HA and (v) flavanone base; 2O; O-Pen; O-Hex; HA (3.4 mg and 3.0 mg, respectively).

### NMR condition

The samples were dissolved in CD_3_OD containing tetramethylsilane (TMS) as an internal standard and analyzed using an NMR spectrometer (JNM-ECA500; JEOL, Japan). Data processing was performed using the JEOL Delta software. ^1^H NMR, ^13^C NMR, correlation spectroscopy (COSY), heteronuclear single quantum correlation (HSQC), heteronuclear multiple quantum correlation (HMBC), distortionless enhancement by polarization transfer (DEPT), and total correlation spectroscopy (TOCSY) spectra were obtained. Chemical shifts were referenced to TMS and subsequently assigned.

### Elucidation of the predicted biosynthetic pathway

Authentic standards of 6”-O-malonyldaidzin (≥90.0% purity by HPLC; Fujifilm Wako Pure Chemical Corporation, Japan) and 1-piperideine (Angene International Limited) were dissolved in 50% MeOH to a final concentration of 500 μM. The two solutions were mixed in equal volumes (1:1) and incubated overnight at either room temperature or in a water bath maintained at 37 °C. After incubation, the reaction mixtures were analyzed by LC-MS/MS.

## Results

### A workflow for data processing and curations of ^12^C and ^13^C paired LC-MS data

We first estimated the carbon numbers of the licorice metabolite peaks using the isotope-labeled tracking function implemented in MS-DIAL 4 [20] (Fig. 1A). The paired LC-MS/MS data from labeled and unlabeled samples were analyzed for each tissue of each plant species, where the labeled samples contained 97 atom% ^13^C, as certified by the supplier. Therefore, all carbon atoms in the metabolites derived from the labeled samples were assumed to be fully substituted with the stable isotope ^13^C. Based on this assumption, the carbon numbers were estimated from the *m/z* shifts observed between the labeled and unlabeled ions in the same or mostly similar retention time region. After the MS-DIAL automatic assignment of ^12^C and ^13^C peak pairs between the two LC-MS data files, the paired results were manually curated by confirming similarities in retention times and chromatographic peak shapes, as well as confirming metabolite ion redundancies. In untargeted metabolomics data using LC-MS/MS, many unannotated ions would originate from contaminants, artifacts, in-source fragment ions, or unexpected ion forms of known metabolites [22,23]. We used the MS-DIAL peak character estimator function, suggesting cluster ions and in-source fragment ions, in addition to the annotation of peaks with similar chromatographic profiles [20]. The remaining ions were defined as “presumed metabolite ion features,” whose adduct forms were mostly estimated as [M+H]^+^, [M+Na]^+^, [M-H]^-^, or [M+HCOO]^-^. Molecular formula estimation for the presumed metabolite ion features was performed using MS-FINDER [21], incorporating the carbon number information derived from the ^13^C labeling experiments (Fig. 1B), where the mass tolerances were set to 5 ppm for MS1 and 10 ppm for MS2. The MS/MS spectral similarity between metabolic peaks with different precursor *m/z* values was calculated using the Bonanza score [14,24] for all presumed molecular ion features. Each peak in the MS/MS spectrum was treated as a node, and edges were created between pairs of nodes showing similarity scores above a defined threshold to construct a molecular network in the Cytoscape environment [25] (Fig. 1C).

In an MS/MS spectral similarity-based molecular network, nodes connected by edges with high spectral similarity can be hypothesized to have the same substructure motifs. Using this assumption, the MS/MS spectra of unknown ions were characterized with prior knowledge of diagnostic product ions or neutral loss features representing substructure motifs, such as flavonoid aglycons and glycosides (**Supplementary Data 1**), with the estimated carbon number and molecular formula information (Fig. 1D).

Candidate molecular formulas were first narrowed down using the carbon number, and the final molecular formula was estimated using MS-FINDER. (C) MS/MS spectral similarity calculation. Similarity was evaluated using the Bonanza score (≥ 0.8). (D) Structural estimation of unknown ions using MS/MS data. Each ion was represented as a node, and edges were formed between nodes showing spectral similarity to constructing MS/MS networks. The structures were estimated based on information from known compounds, predicted molecular formulas, and MS/MS spectral knowledge.

### Results of metabolite annotations using molecular network with isotope-labeled plants

Hydrophilic fractions were extracted from the roots and leaves/stems of both labeled and unlabeled *G. uralensis* and *G. glabra*, followed by reverse-phase LC-MS/MS (**Supplementary Data 2**). In the four unlabeled samples, the number of peaks with MS/MS information was 7,753 in positive ion mode and 5,825 in negative ion mode. By searching mass spectral libraries, 1,304 ions in the positive ion mode and 714 ions in the negative ion mode were annotated with a dot product threshold of 80% (**Supplementary Data 2**), indicating that approximately 15% of the MS/MS acquired peaks were characterized. After carbon number estimation and redundant feature reduction, 1,636 and 1,424 presumed metabolite ion features were obtained in the positive and negative ion modes, respectively (**Supplementary Data 3** and **Supplementary Fig. 1**). Molecular formula predictions with carbon number information, followed by a search for structure candidates, were performed using MS-FINDER [21].

For each species and tissue, MS/MS spectral similarity-based molecular networks were constructed using both the presumed metabolite ion features and authentic standard spectral records from the MassBank of North America (MoNA) and NIST 20 spectral libraries (Fig. 2A and **Supplementary Fig. 2**). In this study, the assignment of substructure information for unknown peaks was performed by integrating knowledge of the carbon number, formula, diagnostic product ion, diagnostic neutral loss, and node/edge connectivity in the molecular network. The newly propagated metabolic ion structures primarily belong to flavonoids and triterpenoids. Based on previously defined criteria [14], peaks annotated by the MS/MS spectral library search were categorized as annotation level 2.1, whereas peaks putatively annotated by integrating the above information were classified as level 2.2. The peaks for which molecular formulas were assigned were categorized as level 3.1, and those for which only the carbon numbers were determined were classified as level 3.2. Peaks with annotation levels of 2.1 or 2.2 were considered structurally annotated metabolites. The number of level 2.2 metabolites contained 48 (root) and 8 (leaf/stem) ions in *G. uralensis* and 34 (root) and 16 (leaf/stem) ions in *G. glabra* in the positive ion mode, 36 (root) and 10 (leaf/stem) ions in *G. uralensis*, and 30 (root) and 12 (leaf/stem) ions in *G. glabra* in the negative ion mode (Fig. 2B). With peaks annotated by spectral library-based spectral matching (level 2.1), 555 ions in the positive ion mode and 460 ions in the negative ion mode were regarded as structurally annotated, indicating that approximately 33% of the putative metabolite-ion features were structurally characterized. In addition, the number of peaks classified as level 3.1 (molecular formula assigned by MS-FINDER [21]) was 409 and 44 ions in the positive and negative ion modes, respectively. When annotation levels 2.1, 2.2, and 3.1 were combined, molecular formula information was assigned to approximately 48% of the putative metabolite ion features (**Supplementary Data 3**). The annotated ions were classified into the “class” layer of ClassyFire program [26] (Fig. 2C), indicating that flavonoids are the most annotated metabolite class, followed by prenol lipids, including triterpenoids. Prenol lipids and isoflavonoids were predominantly enriched in the roots, whereas stilbenes were mainly enriched in the leaves and stems. Previous studies have reported that isoflavonoids and triterpenoids accumulate mainly in the roots of licorice [27], whereas several stilbene derivatives have been isolated from the aerial parts of *Glycyrrhiza* spp. [28]. These observations are consistent with our results, suggesting tissue-specific metabolite localization in licorice.

**Figure 2.**
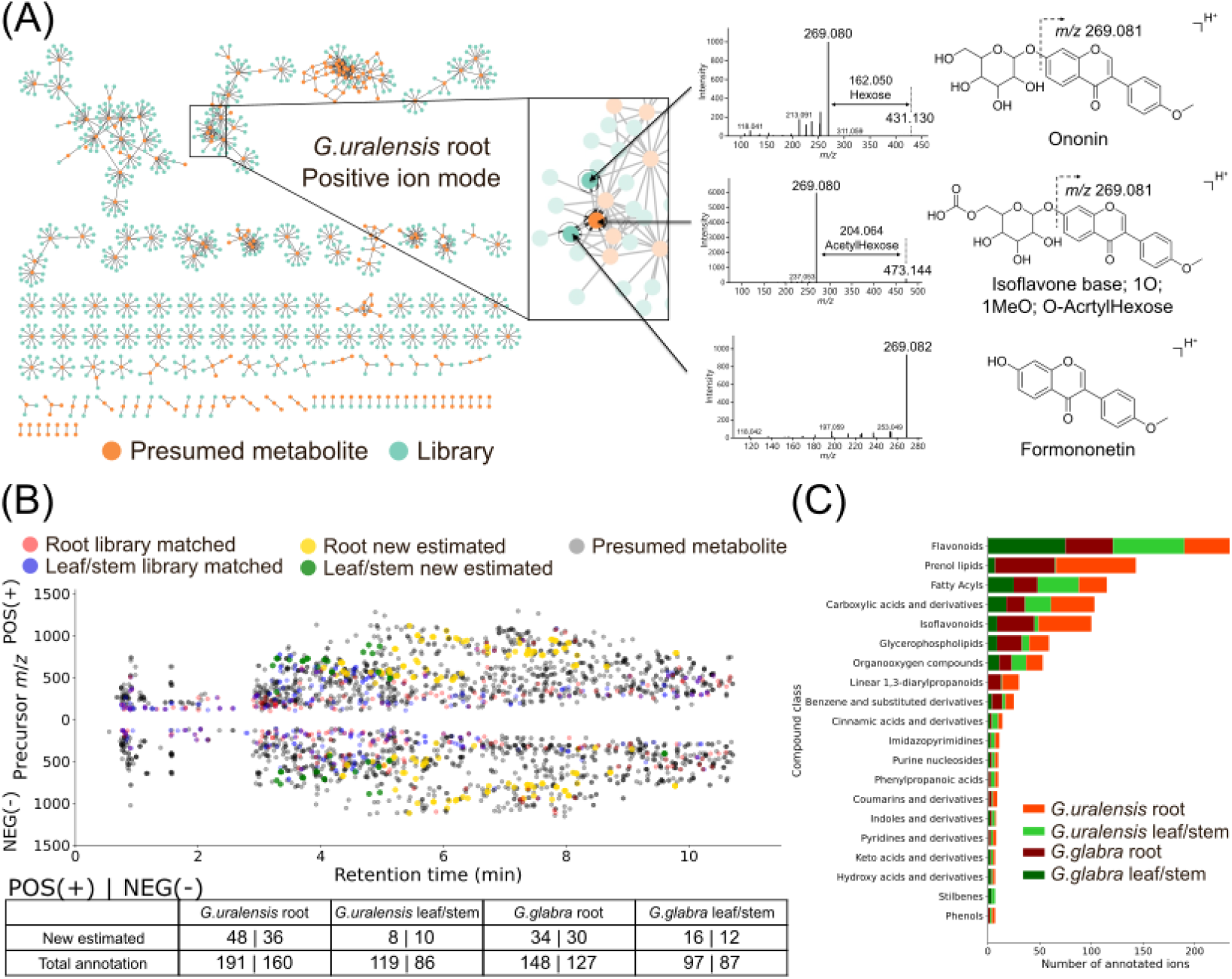
Chemical characterizations to the metabolic ion features detected in *Glycyrrhiza* samples. (A) MS/MS spectral similarity–based molecular networks were constructed using both the presumed metabolite ion features and authentic standard spectral records from spectral libraries. The figure shows the result of *G. uralensis* root in positive ion mode. The three spectra share a common product ion at *m/z* 269.081 (isoflavone base; 1O; 1MeO), and the structures of the presumed metabolite ion was inferred from the unique neutral loss feature. (B) Scatter plot of metabolite-derived peaks. Annotation level 2.1 peaks were mapped in red (root) and blue (leaf/stem), while annotation level 2.2 peaks were mapped in yellow (root) and green (leaf/stem). The peaks assigned to annotation levels 3.1 and 3.2 are mapped in gray. (C) Compound classes of annotated ions (top 20). The ions derived from each sample were distinguished by different colors.

### Overview of the licorice metabolite diversity and discovery of root-specific novel alkaloids

To clarify the differences in metabolite localization among tissues, we described the peak abundance ratios in the nodes of the molecular spectrum network using all putative metabolite ion features (Fig. 3A). Triterpenoids formed a distinct cluster (labeled as “a” in Fig. 3A) with a high proportion of root-derived ions. This observation is consistent with previous reports showing that genes involved in triterpenoid biosynthesis are highly expressed in roots [27,29]. Glycyrrhizin, a triterpenoid found in licorice, contributes to the regulation of soil microbial communities [30]. Therefore, root-specific accumulation of triterpenoids is consistent with their proposed physiological roles. On the other hand, the metabolite class of stilbenes was enriched in leaves/stems represented as the cluster label “b.” Stilbenes are secondary defense metabolites (phytoalexins) induced by ultraviolet radiation [31]. Several stilbenes have been specifically detected in the aerial parts of *Glycyrrhiza* spp. [28], and their distribution is consistent with their proposed physiological functions.

**Figure 3.**
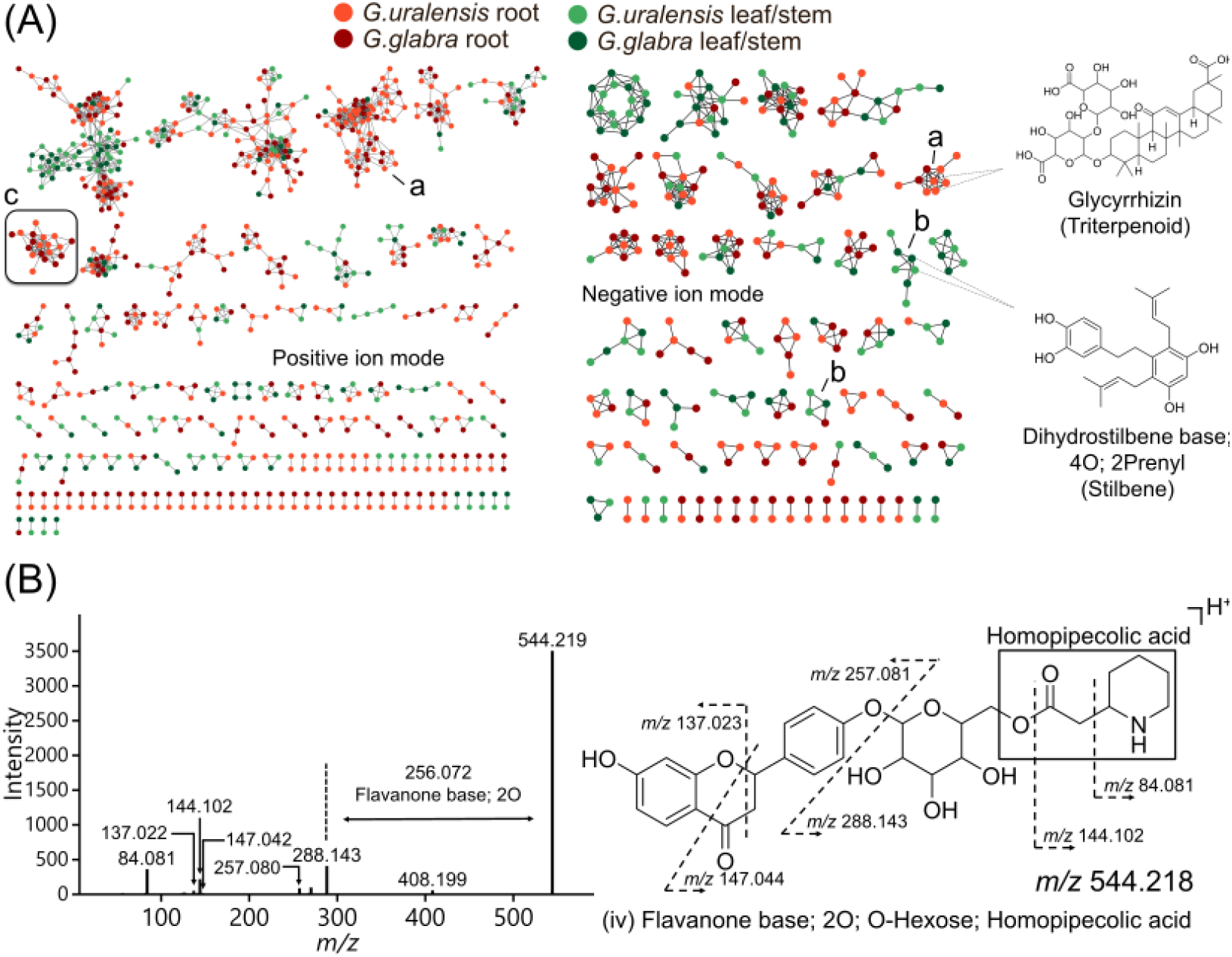
Molecular spectral network for carbon-determined metabolite ion features, discovering novel alkaloids in the root of *Glycyrrhiza* species. (A) MS/MS network constructed among data-cleanup ions, with node colors indicating the sample origin of each ion. Several tissue-specific metabolites were identified. (B) MS/MS spectrum of a root-specific novel alkaloid and corresponding fragment assignments. The unique fragment ions were detected at *m/z* 288.143, 144.102, and 84.081.

Furthermore, we focused on a molecular cluster (labeled as “c”) from the positive ion mode data, which only contained unknown ions detected in the root samples (Fig. 3A). The predicted molecular formulae of these unknown ions contained nitrogen atoms, suggesting that they are alkaloid-like metabolites. Most of the unknown ions within this cluster shared characteristic fragment ions at *m/z* 288.143, 144.102, and 84.081. Similar fragment ion patterns have also been reported in the mass spectrometric analysis of *Ononis* spp., members of the Fabaceae family, and were reported to originate from homopipecolic acid-derived fragments [32,33] (Fig. 3B). In plants, the occurrence of monosubstituted piperidine derivatives is rare, and our study provides the first evidence of their presence in *Glycyrrhiza* spp. To exclude the possibility that these metabolites were sample-specific artifacts, eight commercially available licorice products were analyzed. Nitrogen-containing metabolites with similar MS/MS patterns were detected in all samples (**Supplementary Fig. 3**), indicating that these compounds were widely distributed in licorice roots. Hereafter, the homopipecolic acid (HA) substructure is used as a representative structural description of unknown alkaloids. In this study, five homopipecolic acid-containing metabolites were detected, which contains (i) isoflavone base; O; MeO; O-Hex; HA, (ii) isoflavone base; O; 2MeO; O-Hex; HA, (iii) isoflavone base; 2O; MeO; O-Hex; HA, (iv) flavanone base; 2O; O-Hex; HA, and (v) flavanone base; 2O; O-Pen; O-Hex; HA. Compounds (iii), (iv), and (v) have not been previously reported in either *Ononis* spp. or *Glycyrrhiza* spp. and are therefore presumed to be entirely novel at the whole-structure level.

Among the homopipecolic acid–containing metabolites estimated in this study, compounds (iv) and (v) were considered to be relatively abundant in licorice. Therefore, these compounds were purified and isolated from commercially available licorice samples (Tohoku, Uchida) and subjected to structural elucidation using nuclear magnetic resonance (NMR) spectroscopy. NMR experiments, including ^1^H NMR, ^13^C NMR, HSQC, HMBC, COSY, and TOCSY were conducted. Although the complete separation of compounds (iv) and (v) by HPLC was not possible (**Fig. 4A**), two fractions with different abundance ratios of compound (v) were obtained. By exploiting these differences in relative abundance, the corresponding signals were distinguished, and the overall structures of both compounds were estimated using NMR spectroscopy (**Fig. 4B**). With respect to the monosubstituted piperidine moiety, multiple methylene and methine proton signals were observed in the aliphatic region of the ^1^H NMR spectrum (**Supplementary Fig. 4**). TOCSY correlations established a continuous spin system from H-17 to H-15, consistent with a six-membered ring structure (**Supplementary Fig.5C, D**). Notably, the methine proton at δ_H_ 3.46 (H-11) and the diastereotopic methylene protons at δ_H_ 2.98 and 3.36 (H-15a,b) appeared relatively downfield, suggesting that these protons are adjacent to nitrogen. The corresponding carbons at δ_C_ 54.7 and 46.1 also appeared at relatively downfield chemical shifts, supporting their assignment as carbons neighboring the NH functionality (**Supplementary Fig. 4**). Further analysis of the HMBC spectrum allowed the establishment of connectivity between the individual structural units (**Fig. 4B** and **Supplementary Fig. 6**). The linkage position of the pentose (Pen) unit in compound (v) could not be established. However, liquiritin apioside has been reported in licorice, and based on these biosynthetic precedents, the Pen unit is likely attached to the 2′′-OH of the Hex [34]. This study is the first to confirm the presence of homopipecolic acid-containing metabolites in licorice based on the information corresponding to annotation level 1. Based on the observation that homopipecolic acid-containing metabolites have been detected in leguminous plants such as *Ononis* spp. and *Glycyrrhiza* spp., we investigated the presence of homopipecolic acid in *G. max*, a representative legume species. Homopipecolic acid-containing metabolites were characterized based on three characteristic fragment ions (*m/z* 288.143, 144.102, and 84.081). A metabolite estimated to be daidzein (isoflavone base; 2O) conjugated with homopipecolic acid and a hexose (isoflavone base; 2O; O-Hex; HA) was discovered in the roots. This study provides the first confirmation of the presence of homopipecolic acid-containing metabolites in *G. max* (**Fig. 4C**).

**Figure 4.**
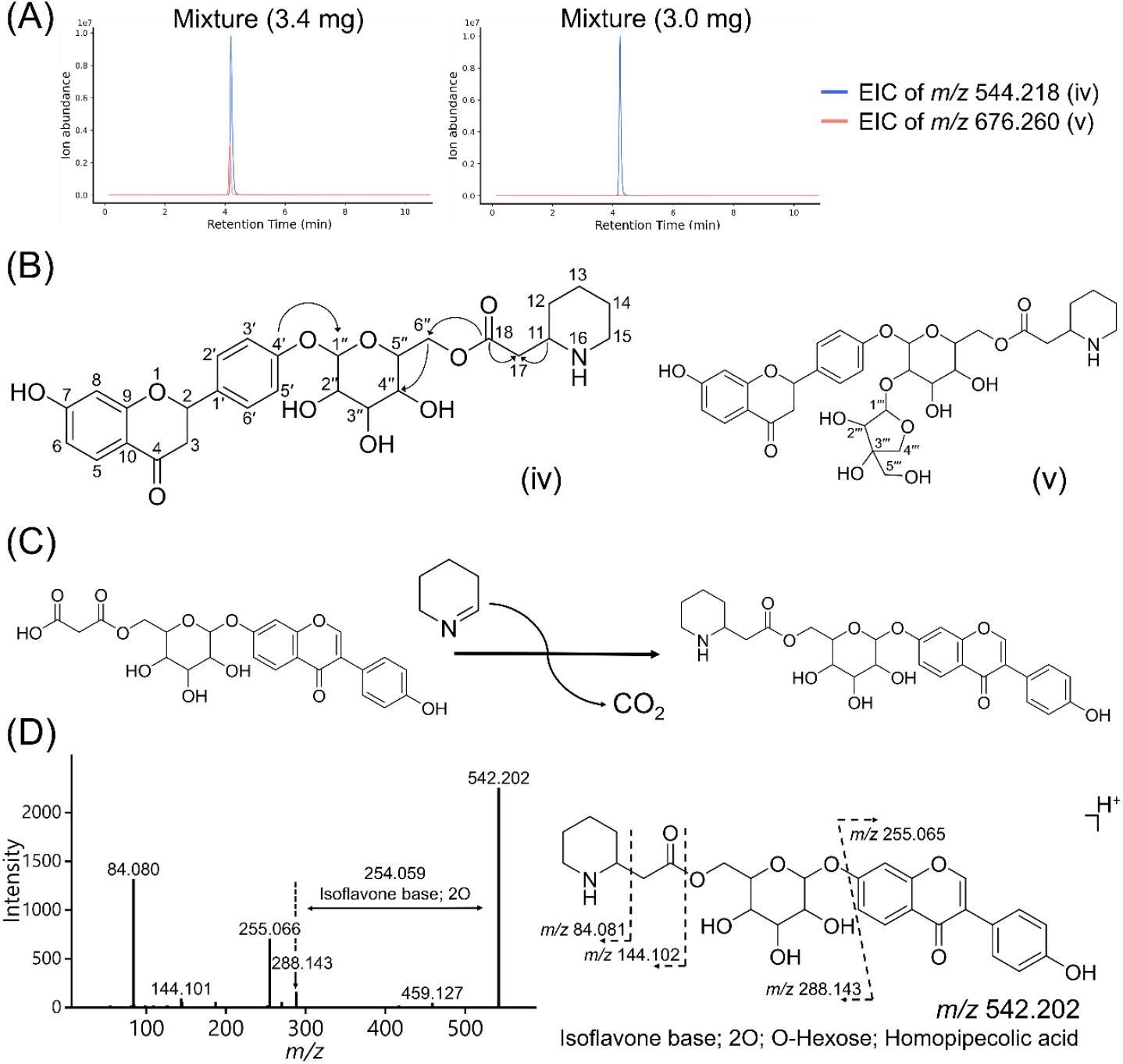
Elucidating licorice-specific alkaloid structures and their potential biochemical pathway. (A) Extracted ion chromatograms corresponding to compounds (iv) (blue) and (v) (red) in the fraction subjected to isolation and purification. (B) Structures of compounds (iv) and (v) proposed based on NMR analysis. Arrows indicate HMBC (^13^C → ^1^H) correlations. (C) A homopipecolic acid– containing metabolite identified in *Glycine max*, with daidzein as the aglycone. Characteristic fragment ions were detected at *m/z* 288.143, 144.102, and 84.081. (D) Proposed biosynthetic pathway of the novel alkaloid. 6”-O-malonyldaidzin reacts with Δ^1^-piperideine to produce a homopipecolic acid-containing metabolite.

Several distinct alkaloid biosynthetic pathways have been identified in legumes. Among them, the quinolizidine alkaloid biosynthetic pathway, which is abundant in *Lupinus* spp., is one of the most famous and well-studied pathways. It is synthesized from lysine via cadaverine and 5-aminopentanal as intermediates [35]. In addition, in ferns of the genus *Lycopodium*, lysine is similarly utilized as a precursor, and Δ^1^-piperideine undergoes a spontaneous non-enzymatic Mannich-like condensation with 3-oxoglutaric acid to form pelletierine, a compound structurally similar to homopipecolic acid [36]. In plants, metabolites often accumulate as flavonoid glycosides conjugated with diverse sugar moieties, which enhance their solubility and stability in vacuoles [37]. Malonyl glucose-containing glycosides have been widely reported in leguminous plants [38]. Given that the malonyl group, like 3-oxoglutaric acid, possesses a β-keto acid moiety, we hypothesized that flavonoid glycosides bearing malonylated glucose could serve as substrates for homopipecolic acid conjugation through a reaction with Δ^1^-piperideine (Fig. 4D). To validate this hypothesis, authentic standards of 6”-O-malonyldaidzin (daidzein [isoflavone base; 2O] + malonyl glucose), which is endogenously detected in *G. max*, and 1-piperideine were mixed and incubated overnight at either room temperature or 37 °C. Fragment ions corresponding to daidzein conjugated with homopipecolic acid and a hexose moiety (isoflavone base; 2O; O-Hex; HA) were detected in both conditions (**Supplementary Fig. 7**). These results suggest that homopipecolic acid is derived from a reaction between 1-piperideine and flavonoid glycosides containing malonyl glucose, and that the final step of this biosynthetic process likely proceeds via a non-enzymatic reaction. Consistent with this hypothesis, potential precursor flavonoid glycosides have been detected in planta, including daidzein conjugated with malonyl glucose (isoflavone base; 2O; O-MHex) in *G. max*, and flavanone base; 2O; O-MHex as well as flavanone base; 2O; O-Pen; O-MHex in *Glycyrrhiza spp*. Notably, in *Glycyrrhiza* spp., these precursor candidates were specifically detected in the roots, which aligned well with the root-specific accumulation of homopipecolic acid-containing metabolites (**Supplementary Fig. 8**).

## Discussion

In the present study, we investigated the diversity of plant-specialized metabolites in *Glycyrrhiza* by combining a molecular networking approach with stable isotope-labeled plants. As a result, 3,060 metabolite-derived peaks were detected and structural annotations were assigned to 1,015 ions, corresponding to approximately 33% of the detected metabolite-derived peaks. Furthermore, 194 of these ions were not registered in the existing and well-known metabolite database KNApSAcK [3]. Among these putative novel metabolites, five homopipecolic acid-containing metabolites were specifically localized in the roots. Of these, two compounds were considered likely to possess novel structures. These two compounds were purified and isolated, and their structures were determined using NMR. Two novel alkaloids were discovered in this study, both of which were confirmed for the first time in *Glycyrrhiza* spp.

Although reports of 1-substituted piperidine derivatives, such as homopipecolic acid, in plants are limited, our study demonstrates that this compound is present not only in *Ononis* spp. but also in *Glycyrrhiza* spp. and *G. max*. All these species belong to the Fabaceae family, and this metabolite has been detected exclusively in the roots of Fabaceae plants. These observations suggested that this metabolite may have been acquired during a lineage-specific evolutionary process in the Fabaceae family. Our results suggest that flavonoid glycosides containing malonyl glucose, which have been widely reported in Fabaceae plants, together with 1-piperideine, may be involved in the final step of the biosynthesis of this metabolite. 1-Piperideine is synthesized from lysine via a pathway involving lysine decarboxylase (LDC) and copper amine oxidase (CuAO) (**Supplementary Fig. 9**). This route has also been reported for the biosynthesis of quinolizidine alkaloids, which are representative alkaloids in Fabaceae plants [35]. In *G. max*, the presence of both LDC and CuAO has been suggested. LDC exhibits particularly high activity in the roots, and CuAO has been reported to show root-specific expression [39,40]. These findings are consistent with the root-specific distribution of metabolites observed in this study. In contrast, metabolites containing homopipecolic acid have not yet been reported in quinolizidine alkaloid-producing plants, such as *Lupinus* spp. Taken together, these results indicate that further analysis of genes related to 1-piperideine metabolism in *Glycyrrhiza* spp. is likely to provide new insights into alkaloid biosynthetic pathways in Fabaceae plants, as well as Fabaceae-specific evolutionary processes.

Homopipecolic acid is a nitrogen-containing compound that exhibits structural similarities to methylphenidate, a central nervous system stimulant. This structural similarity raises the possibility that metabolites containing homopipecolic acid exert physiological effects similar to those of methylphenidates. Both licorice extract and methylphenidate have been reported to increase dopamine and noradrenaline levels [41,42]. Additionally, licorice extracts possess nootropic and anti-amnesic properties [43,44]. These findings suggest that homopipecolic acid–containing metabolites may contribute to the neurological effects of licorice. Plant-derived alkaloids produced in plants are a class of compounds that have long been utilized by humans because of their characteristic chemical properties, and many have been used as pharmaceuticals for the treatment of a wide range of diseases. However, many alkaloids are toxic to humans, herbivores, and insects [43], and this toxicity is considered a chemical defense mechanism [35]. In the present study, we identified previously unreported homopipecolic acid-containing alkaloids that are specifically localized in licorice roots. Further elucidation of their botanical significance and physiological roles may promote the discovery of novel pharmacological activities and contribute to the development of efficient production strategies for licorice.

## Data availability

All raw MS data are available in the MB-POST repository (https://repository.massbank.jp/) under index number MPST000161.0. The original tables and source codes for the Figures displayed in this study are available as Source Data.

## Code availability

MS-DIAL source code is available at https://github.com/systemsomicslab/MsdialWorkbench.

## Acknowledgments

This research was supported by the Japan Agency for Medical Research and Development (AMED) under Infectious Diseases Research and Infrastructure (JP25wm0325071, H.T.), the Japan Science and Technology Agency (JST) Exploratory Research for Advanced Technology (ERATO) (JPMJER2101 to H. T.), JST FOREST program (JPMJFR230H to H.T.), JST NBDC (JPMJND2305 to H.T.), and the JSPS KAKENHI (24K02011, 24H00043, 24H00392, 24K21269, 25H01425, and 25H01426 to H.T.).

## Author Contributions

H.T. designed the study. K.S., Y.T., and Y.M. performed LC-MS/MS experiments and data analyses. K.S., Y.T., S.N., K.N., T.K., and M.Y.H. performed metabolite purifications and NMR analyses. T.M. and A.R. performed biological sample management, and K.S., A.R., and H.T. discussed the genomic interpretations. K.S. and H.T. wrote the manuscript, and all authors contributed to improve the draft. All authors have thoroughly discussed this project and helped improve the manuscript.

## Competing interests

All authors declare no competing interests.

## Notes

### Competing Interest Statement

The authors have declared no competing interest.

## References

[1] Shadma Wahab et al. Glycyrrhiza glabra (Licorice): A Comprehensive Review on Its Phytochemistry, Biological Activities, Clinical Evidence and Toxicology. Plants, 2021 Dec; 10(12): 2751.

[2] Liwei Wu et al. Glycyrrhiza, a commonly used medicinal herb: Review of species classification, pharmacology, active ingredient biosynthesis, and synthetic biology. Journal of Advanced Research, 75 (2025) 249–270.

[3] Farit Mochamad Afendi et al. KNApSAcK family databases: integrated metabolite-plant species databases for multifaceted plant research. Plant Cell Physiol, 2012 Feb;53(2).

[4] Yuping Li et al. Integrated metabolomic and transcriptomic analysis provides insights into the flavonoid formation in different Glycyrrhiza species. Industrial Crops and Products, Volume 208, February 2024, 117796.

[5] Jianping Zhao et al. Metabolite variation and discrimination of five licorice (Glycyrrhiza) species: HPTLC and NMR explorations. Journal of Pharmaceutical and Biomedical Analysis, 220 (2022) 115012.

[6] Chunsheng Xiao et al. Transport of secondary metabolites in plants: Mechanistic insights and transporter engineering for crop improvement. Plant Communications, 6, 101536, December 8 2025.

[7] Lu Yao et al. Subcellular compartmentalization in the biosynthesis and engineering of plant natural products. Biotechnology Advances, 69 (2023) 108258.

[8] Meng-Ling Shih et al. Metabolic flux analysis of secondary metabolism in plants. Metabolic Engineering Communications, 10 (2020) e00123.

[9] Mahboubeh Irani et al. Leaves Antimicrobial Activity of Glycyrrhiza glabra L. Iran J Pharm Res, 2010 Fall;9(4):425–8.

[10] Luca Frattaruolo et al. Antioxidant and Anti-Inflammatory Activities of Flavanones from Glycyrrhiza glabra L. (licorice) Leaf Phytocomplexes: Identification of Licoflavanone as a Modulator of NF-kB/MAPK Pathway. Antioxidants, 2019, 8, 186.

[11] Giovanni Rizzato et al. A new exploration of licorice metabolome. Food Chemistry, 221 (2017) 959–968.

[12] Hiroshi Tsugawa. et al. Metabolomics and complementary techniques to investigate the plant phytochemical cosmos. Natural Product Reports, Volume 38, Issue 10, 2021, Pages 1729–1759.

[13] Xinyu Yuan et al. Analysis of plant metabolomics data using identification-free approaches. Appl Plant Sci, 2025 Mar 1;13(4):e70001.

[14] Hiroshi Tsugawa. et al. A cheminformatics approach to characterize metabolomes in stable isotope-labeled organisms. Nature methods, 16, 295–298(2019).

[15] Patrick Giavalisco et al. High-Resolution Direct Infusion-Based Mass Spectrometry in Combination with Whole 13C Metabolome Isotope Labeling Allows Unambiguous Assignment of Chemical Sum Formulas. Anal. Chem, 2008, 80, 24, 9417–9425.

[16] Patrick Giavalisco et al. 13C Isotope-Labeled Metabolomes Allowing for Improved Compound Annotation and Relative Quantification in Liquid Chromatography-Mass Spectrometry-based Metabolomic Research. Anal. Chem, 2009, 81, 15, 6546–6551.

[17] Asja Ćeranić et al. Preparation of uniformly labelled 13C- and 15N-plants using customised growth chambers. Plant Methods, (2020) 16:46.

[18] Achuthanunni Chokkathukalam et al. Stable isotope-labeling studies in metabolomics: new insights into structure and dynamics of metabolic networks. Bioanalysis, 2014 Feb;6(4):511–524.

[19] Nirin Udomsom et al. Function of AP2/ERF Transcription Factors Involved in the Regulation of Specialized Metabolism in Ophiorrhiza pumila Revealed by Transcriptomics and Metabolomics. Front. Plant Sci., 7, 1–14 (2016).

[20] Hiroshi Tsugawa. et al. A lipidome atlas in MS-DIAL 4. Nature Biotechnology, 38, 1159–1163(2020).

[21] Hiroshi Tsugawa. et al. Hydrogen Rearrangement Rules: Computational MS/MS Fragmentation and Structure Elucidation Using MS-FINDER Software. Analytical Chemistry, 88, 7946 (2016).

[22] Nathaniel G. Mahieu et al. Systems-Level Annotation of a Metabolomics Data Set Reduces 25,000 Features to Fewer than 1,000 Unique Metabolites. Anal Chem., 2017 Oct 3; 89(19): 10397–10406.

[23] Jun Xie et al. Spontaneous In-Source Fragmentation Reaction Mechanism and Highly Sensitive Analysis of Dicofol by Electrospray Ionization Mass Spectrometry. Molecules, 2023 May; 28(9): 3765.

[24] Jayson A Falkner et al. A spectral clustering approach to MS/MS identification of posttranslational modifications. J Proteome Res., 2008 Nov;7(11):4614–22.

[25] Paul Shannon et al. Cytoscape: a software environment for integrated models of biomolecular interaction networks. Genome Res, 2003 Nov; 13(11): 2498–2504.

[26] Yannick Djoumbou Feunang et al. ClassyFire: automated chemical classification with a comprehensive, computable taxonomy. J Cheminform, 2016 Nov 4:8:61.

[27] Kaiqiang Yu et al. Integrative metabolome and transcriptome analyses reveal the differences in flavonoid and terpenoid synthesis between Glycyrrhiza uralensis (licorice) leaves and roots. Food Science and Biotechnology, 2023 Nov 20;33(1):91–101.

[28] T K Lim. Glycyrrhiza glabra. Edible Medicinal and Non-Medicinal Plants, 2015 Oct 22:354–457.

[29] Hikaru Seki et al. Licorice β-amyrin 11-oxidase, a cytochrome P450 with a key role in the biosynthesis of the triterpene sweetener glycyrrhizin. Proc Natl Acad Sci U S A, 2008 Sep 8;105(37):14204–14209.

[30] Masaru Nakayasu et al. Triterpenoid and Steroidal Saponins Differentially Influence Soil Bacterial Genera. Plants, 2021, 10, 2189.

[31] Julie Chong et al. Metabolism and roles of stilbenes in plants. Plant Science, 177 (2009) 143–155.

[32] Nóra Gampe et al. Separation and characterization of homopipecolic acid isoflavonoid ester derivatives isolated from Ononis spinosa L. root. J Chromatogr B Analyt Technol Biomed Life Sci., 2018 Aug 1:1091:21–28.

[33] Nóra Gampe et al. Phytochemical analysis of Ononis arvensis L. by liquid chromatography coupled with mass spectrometry. J Mass Spectrom., 2019 Feb;54(2):121–133.

[34] Hao-Tian Wang et al. Insights into the missing apiosylation step in flavonoid apiosides biosynthesis of Leguminosae plants. Nat Commun, 2023 Oct 20;14:6658.

[35] Astrid Ramírez-Betancourt et al. Unraveling the Biosynthesis of Quinolizidine Alkaloids Using the Genetic and Chemical Diversity of Mexican Lupins. Diversity, 2021, 13, 375.

[36] Juan Wang et al. Deciphering the Biosynthetic Mechanism of Pelletierine in Lycopodium Alkaloid Biosynthesis. Organic Letters, 2020, 22, 21, 8725–8729.

[37] Julien Le Roy et al. Glycosylation Is a Major Regulator of Phenylpropanoid Availability and Biological Activity in Plants. Front Plant Sci., 2016 May 26;7:735.

[38] Muhammad Z Ahmad et al. Isoflavone Malonyltransferases GmIMaT1 and GmIMaT3 Differently Modify Isoflavone Glucosides in Soybean (Glycine max) under Various Stresses. Front Plant Sci., 2017 May 16:8:735.

[39] M. Ohe et al. Putative occurrence of lysine decarboxylase isoforms in soybean (Glycine max) seedlings. Amino Acids, 2009 Jan;36(1):65–70

[40] Costas Delis et al. A root- and hypocotyl-specific gene coding for copper-containing amine oxidase is related to cell expansion in soybean seedlings. Journal of Experimental Botany, Volume 57, Issue 1, January 2006, Pages 101–111.

[41] Rafał R. Jaeschke et al. Methylphenidate for attention-deficit/hyperactivity disorder in adults: a narrative review. Psychopharmacology, 2021; 238(10): 2667–2691.

[42] Haidy E. Michel et al. Prepulse inhibition (PPI) disrupting effects of Glycyrrhiza glabra extract in mice: A possible role of monoamines. Neuroscience Letters, Volume 544, 7 June 2013, Pages 110–114.

[43] Kosuri Kalyan Chakravarthi et al. Beneficial effect of aqueous root extract of Glycyrrhiza glabra on learning and memory using different behavioral models: An experimental study. J Nat Sci Biol Med, 2013 Jul-Dec;4(2):420–425.

[44] Dinesh Dhingra et al. Memory enhancing activity of Glycyrrhiza glabra in mice. Journal of Ethnopharmacology, 91 (2004) 361–365.

