## Supplementary material for "Stable isotope–assisted computational mass spectrometry reveals root-specific alkaloids in *Glycyrrhiza* species": Source data: 20250728_Scatter.ipynb

```
# %% [code]
import matplotlib.pyplot as plt
import pandas as pd

# CSVファイルからデータを読み込む
df_all_g = pd.read_csv('D:/20221024_Glycyrrhiza_sawai/Scatter plot/20251201/glabra/glabra_all.csv')
df_all_u = pd.read_csv('D:/20221024_Glycyrrhiza_sawai/Scatter plot/20251201/uralensis/uralensis_all.csv')

df_g = pd.read_csv('D:/20221024_Glycyrrhiza_sawai/Scatter plot/20251201/glabra/glabra_clean-up.csv')
df_u= pd.read_csv('D:/20221024_Glycyrrhiza_sawai/Scatter plot/20251201/uralensis/uralensis_clean-up.csv')

# %% [code]
import matplotlib.pyplot as plt
import pandas as pd

# 色・サイズ・透明度設定
peaks_g_u = {
    'Unknown': {'color': 'grey', 'alpha': 0.3, 'size': 20},
    'Data_clean-up': {'color': 'black', 'alpha': 0.5, 'size': 20},
    'uralensis_root_Library-matched': {'color': 'red', 'alpha': 0.5, 'size': 35},
    'glabra_root_Library-matched': {'color': 'red', 'alpha': 0.5, 'size': 35},
    'uralensis_leaf_Library-matched': {'color': 'blue', 'alpha': 0.5, 'size': 35},
    'glabra_leaf_Library-matched': {'color': 'blue', 'alpha': 0.5, 'size': 35}
}

# --- プロット作成 ---
fig, ax = plt.subplots(figsize=(15, 6))

# 2つのデータセットをループ
for df_all, label_prefix in zip([df_all_g, df_all_u], ['glabra_all', 'uralensis_all']):
    
    # Annotation_tag ごとにプロット
    for group, props in peaks_g_u.items():

        data = df_all[df_all['Annotation_tag'] == group]

        if data.empty:
            continue

        ax.scatter(
            data['RT (min)'],
            data['Precursor m/z'],
            label=f"{label_prefix}_{group}",
            color=props['color'],
            alpha=props['alpha'],
            edgecolors=props['color'],
            s=props['size']
        )

# --- 軸設定 ---
ax.set_xlabel('Retention time (min)', size=20)
ax.set_ylabel('Precursor $m/z$', size=20)
ax.set_xlim(0, 11.5)
ax.set_ylim(-1500, 1550)

yticks = [-1500, -1000, -500, 0, 500, 1000, 1500]
ax.set_yticks(yticks)
ax.set_yticklabels([abs(y) for y in yticks])

ax.tick_params(axis='both', labelsize=15)

# 不要な枠線を消す
ax.spines['right'].set_visible(False)
ax.spines['top'].set_visible(False)

plt.tight_layout()
#plt.legend(bbox_to_anchor=(1.05, 1), loc='upper left')
plt.savefig("D:/20221024_Glycyrrhiza_sawai/Scatter plot/20251201/Scatter_Plot_all.svg", dpi=300)
plt.show()


# %% [code]
import matplotlib.pyplot as plt
import pandas as pd


# new-annotationに対する色・透明度・サイズ設定
annotation_props = {
    'Data_clean-up':                 {'color': 'black',  'alpha': 0.3, 'size': 30},
    'uralensis_root_Library-matched':      {'color': 'red',    'alpha': 0.3, 'size': 30},
    'glabra_root_Library-matched':         {'color': 'red',    'alpha': 0.3, 'size': 30},
    'uralensis_leaf_Library-matched':      {'color': 'blue',   'alpha': 0.3, 'size': 30},
    'glabra_leaf_Library-matched':         {'color': 'blue',   'alpha': 0.3, 'size': 30},
    'uralensis_root_New-estimation':  {'color': 'gold', 'alpha': 0.75, 'size': 50},    
    'glabra_root_New-estimation':     {'color': 'gold', 'alpha': 0.75, 'size': 50},
    'uralensis_leaf_New-estimation':  {'color': 'green',  'alpha': 0.75, 'size': 50},
    'glabra_leaf_New-estimation':     {'color': 'green',  'alpha': 0.75, 'size': 50},
}   

fig, ax = plt.subplots(figsize=(15,6))

# df_g, df_u 両方まとめて処理
for df, label_prefix in zip([df_g, df_u], ['glabra', 'uralensis']):
    for annotation, props in annotation_props.items():
        # そのannotationのデータがあるかチェック
        if annotation in df['Annotation_tag'].values:
            subset = df[df['Annotation_tag'] == annotation]
            ax.scatter(
                subset['RT (min)'],
                subset['Precursor m/z'],
                label=f"{label_prefix}_{annotation}",
                color=props['color'],
                alpha=props['alpha'],
                edgecolors=props['color'],
                s=props['size']
            )

# 軸設定
ax.set_xlabel('Retention time (min)', size=25)
ax.set_ylabel('Precursor $m/z$', size=25)
ax.set_xlim(0, 11.5)
ax.set_ylim(-1500, 1550)

# y軸目盛りのラベル調整（例）
yticks = [-1500, -1000, -500, 0, 500, 1000, 1500]
ax.set_yticks(yticks)
ax.set_yticklabels([abs(y) for y in yticks])

ax.tick_params(axis='both', labelsize=20)

# 右上の枠線と上の枠線を非表示に
ax.spines['right'].set_visible(False)
ax.spines['top'].set_visible(False)


plt.tight_layout()
plt.savefig(
    "D:/20221024_Glycyrrhiza_sawai/Scatter plot/20251201/Scatter_Plot_clean-up.svg",
    format="svg"
)
plt.show()
```
