## Supplementary material for "Stable isotope–assisted computational mass spectrometry reveals root-specific alkaloids in *Glycyrrhiza* species": Source data: class.ipynb

```
# %% [code]
import pandas as pd
import matplotlib.pyplot as plt

# =====================
# CSV読み込み
# =====================
df = pd.read_csv("D:/発表関連/原稿/class_counts_per_sample.csv")

# Part列をindexに
df = df.set_index("Part")

# =====================
# データ整形
# =====================
# 転置（compound class を行にする）
df_t = df.T

# 合計値で並び替え
df_t["total"] = df_t.sum(axis=1)
df_t = df_t.sort_values("total", ascending=True)
df_t = df_t.drop(columns="total")

# =====================
# 色設定（画像に近い色）
# =====================
colors = {
    "uralensis_root": "#ff4500",   # オレンジ
    "uralensis_leaf": "#32cd32",   # 明るい緑
    "glabra_root": "#8b0000",# 濃い赤
    "glabra_leaf": "#006400" # 濃い緑
}

# =====================
# グラフ描画
# =====================
fig, ax = plt.subplots(figsize=(10, 14))

left = pd.Series([0]*len(df_t), index=df_t.index)

for col in df_t.columns:
    ax.barh(
        df_t.index,
        df_t[col],
        left=left,
        color=colors[col],
        label=col
    )
    left += df_t[col]

# =====================
# ラベル設定
# =====================
ax.set_xlabel("Number of annotated ions", fontsize=16)
ax.set_ylabel("Compound class", fontsize=18)

# 凡例
#ax.legend(title="Tissue", bbox_to_anchor=(1.02, 1), loc="upper left")

# レイアウト
plt.tight_layout()

# 保存（任意）
# plt.savefig("compound_class_distribution.png", dpi=300)

plt.show()

# %% [code]
import pandas as pd
import matplotlib.pyplot as plt

# =====================
# CSV読み込み
# =====================
df = pd.read_csv("D:/発表関連/原稿/class_counts_per_sample.csv")

# Part列をindexに
df = df.set_index("Part")

# =====================
# データ整形
# =====================
# 転置（compound class を行にする）
df_t = df.T

# 合計値で並び替え
df_t["total"] = df_t.sum(axis=1)
df_t = df_t.sort_values("total", ascending=False)

# 上位15クラスのみ
df_t = df_t.head(20)

# total列削除
df_t = df_t.drop(columns="total")

# 表示のため昇順に（グラフを見やすく）
df_t = df_t[::-1]

# =====================
# 色設定
# =====================
colors = {
    "uralensis_root": "#ff4500",
    "uralensis_leaf": "#32cd32",
    "glabra_root": "#8b0000",
    "glabra_leaf": "#006400"
}

# =====================
# グラフ描画
# =====================
fig, ax = plt.subplots(figsize=(10, 10))

left = pd.Series([0]*len(df_t), index=df_t.index)

for col in df_t.columns:
    ax.barh(
        df_t.index,
        df_t[col],
        left=left,
        color=colors[col],
        label=col
    )
    left += df_t[col]

# =====================
# 軸ラベル
# =====================
ax.set_xlabel("Number of annotated ions", fontsize=17.5)
ax.set_ylabel("Compound class", fontsize=17.5)

ax.tick_params( labelsize=15)

# =====================
# 外枠設定（左と下だけ表示）
# =====================
ax.spines["top"].set_visible(False)
ax.spines["right"].set_visible(False)
ax.spines["left"].set_visible(True)
ax.spines["bottom"].set_visible(True)

# tickだけ残す
ax.tick_params(left=True, bottom=True)

plt.tight_layout()
plt.savefig("D:/発表関連/原稿/Class.svg", dpi=300)
plt.show()
```
