## Supplementary material for "Stable isotope–assisted computational mass spectrometry reveals root-specific alkaloids in *Glycyrrhiza* species": Source data: Gly4,5.ipynb

```
# %% [code]
import pymzml
import matplotlib.pyplot as plt

run = pymzml.run.Reader("D:/Gly/20250522/Data/20250522_Gly_5_5_POS_1.mzML")

target_mz_1 = 544.2177
target_mz_2 = 676.2599
tolerance = 0.01

times = []
intensities_1 = []
intensities_2 = []

for spectrum in run:
    if spectrum.ms_level == 1:

        rt = spectrum.scan_time_in_minutes()

        intensity_1 = sum(
            inten for mz, inten in spectrum.peaks("raw")
            if abs(mz - target_mz_1) <= tolerance
        )

        intensity_2 = sum(
            inten for mz, inten in spectrum.peaks("raw")
            if abs(mz - target_mz_2) <= tolerance
        )

        times.append(rt)
        intensities_1.append(intensity_1)
        intensities_2.append(intensity_2)

# %% [code]
plt.figure(figsize=(8,4))

plt.plot(
    times,
    intensities_1,
    color='royalblue',
    linewidth=1,
    linestyle='-',
    #label=f"m/z {target_mz_1}"
)

plt.plot(
    times,
    intensities_2,
    color='salmon',
    linewidth=1,
    linestyle='-',
    #label=f"m/z {target_mz_2}"
)

plt.xlabel("Retention Time (min)", fontsize=15)
plt.ylabel("Ion abundance", fontsize=15)
#plt.title("EIC", fontsize=15)

ax = plt.gca()
ax.spines['top'].set_visible(False)
ax.spines['right'].set_visible(False)

ax.spines['left'].set_linewidth(1.5)
ax.spines['bottom'].set_linewidth(1.5)

plt.legend()

plt.savefig("D:/発表関連/原稿/Figure/Gly5.svg", dpi=300)
plt.show()
```
