## Supplementary material for "Stable isotope–assisted computational mass spectrometry reveals root-specific alkaloids in *Glycyrrhiza* species": Source data: chromatogram_Com4.ipynb

```
# %% [code]
import pymzml
import matplotlib.pyplot as plt

# ======================
# ① データ読み込み・保存
# ======================
def extract_eic(files, target_mz, tolerance=0.01):
    eic_data = {}  # {label: (times, intensities)}

    for i, file in enumerate(files):
        run = pymzml.run.Reader(file)

        times, intensities = [], []
        for spectrum in run:
            if spectrum.ms_level == 1:
                rt = spectrum.scan_time_in_minutes()
                intensity = sum(
                    inten for mz, inten in spectrum.peaks("raw")
                    if abs(mz - target_mz) <= tolerance
                )
                times.append(rt)
                intensities.append(intensity)

        eic_data[file] = (times, intensities)

    return eic_data

# ======================
# 使用例
# ======================
files = [    
    "D:/Gly/20231117_8sample/POS/all/20231117_Glycyrrhiza_1_POS_1.mzML",
    "D:/Gly/20231117_8sample/POS/all/20231117_Glycyrrhiza_2_POS_1.mzML",
    "D:/Gly/20231117_8sample/POS/all/20231117_Glycyrrhiza_3_POS_1.mzML",
    "D:/Gly/20231117_8sample/POS/all/20231117_Glycyrrhiza_4_POS_1.mzML",
    "D:/Gly/20231117_8sample/POS/all/20231117_Glycyrrhiza_5_POS_1.mzML",
    "D:/Gly/20231117_8sample/POS/all/20231117_Glycyrrhiza_6_POS_1.mzML",
    "D:/Gly/20231117_8sample/POS/all/20231117_Glycyrrhiza_7_POS_1.mzML",
    "D:/Gly/20231117_8sample/POS/all/20231117_Glycyrrhiza_8_POS_1.mzML"
        ]

# データ抽出
target_mz = 544.218
tolerance = 0.01
eic_data = extract_eic(files, target_mz, tolerance)


# %% [code]
import matplotlib.pyplot as plt
import numpy as np

# ======================
# ② プロット関数
# ======================
def plot_eic(eic_data, labels=None, colors=None,
             offset_x=0.5, offset_y=100000,
             target_mz=None, tolerance=None):

    plt.figure(figsize=(8, 4))

    # y 軸に振る文字
    y_labels = ["a", "b", "c", "d", "e", "f", "g", "h"]

    # 右側に文字を書くために最大RTを取得
    all_times = []
    for times, _ in eic_data.values():
        all_times.extend(times)
    max_rt = max(all_times)

    for i, (file, (times, intensities)) in enumerate(eic_data.items()):
        # サンプルごとに x, y をシフト
        shifted_times = [t + i * offset_x for t in times]
        shifted_intensities = [y + i * offset_y for y in intensities]

        color = colors[i] if colors else "black"

        # クロマト描画
        plt.plot(
            shifted_times,
            shifted_intensities,
            color=color,
            linewidth=1,
        )

        # ===== 各クロマトの右側に a, b, c... を表示 =====
        plt.text(
            max(shifted_times) + 0.2,   # 右側に少し余白
            i * offset_y,               # クロマトの基準y
            y_labels[i],
            fontsize=10,
            va="center",
            ha="left",
                    )

    # 軸ラベル
    plt.xlabel("Retention Time (min)", fontsize=15)
    plt.ylabel("Ion abundance", fontsize=1)

    # ===== y 軸の数値を表示 =====
    ax = plt.gca()
    ax.tick_params(axis="y", labelsize=10)

    # x 軸目盛は消す（論文向け）
    plt.xticks([])

    # タイトル
    if target_mz and tolerance:
        plt.title(
            f"Compound ⅳ (EIC of $\it{{m}}$$\it{{/}}$$\it{{z}}$ "
            f"{target_mz} ± {tolerance})",
            fontsize=15
        )

    # 枠線調整
    ax.spines["top"].set_visible(False)
    ax.spines["right"].set_visible(False)
    ax.spines["left"].set_linewidth(1.2)
    ax.spines["bottom"].set_linewidth(1.2)

    plt.tight_layout()
    plt.savefig("D:/発表関連/原稿/Kanzo.svg", dpi=300)
    plt.show()


# ======================
# 使用例
# ======================

colors = ["black"] * 8

plot_eic(
    eic_data,
    colors=colors,
    offset_x=1,
    offset_y=20000,
    target_mz=target_mz,
    tolerance=tolerance
)
```
