## Supplementary material for "Stable isotope–assisted computational mass spectrometry reveals root-specific alkaloids in *Glycyrrhiza* species": Source data: 2026_Supply9.ipynb

```
# %% [code]
import matplotlib.pyplot as plt
import numpy as np

# データ
labels = ["$G. uralensis$ root", "$G. uralensis$ leaf/stem", "$G. glabra$ root", "$G. glabra$ leaf/stem"]

data_505 = [10373, 32, 51386, 64]
data_637 = [10785, 17, 22403, 57]

x = np.arange(len(labels))

# 図作成
fig, axes = plt.subplots(2, 1, figsize=(6, 8))

# カラー設定（例に近い色）
colors = ["#FF5733", "#4DAA64", "#9E0000", "#005F3C"]

# (A) m/z = 505
axes[0].bar(x, data_505, color=colors)
axes[0].set_title("$m/z$ = 505.134", size = 15)
axes[0].set_ylabel("Peak height", size = 15)
axes[0].set_xticks(x)
axes[0].set_xticklabels(labels)


# (B) m/z = 637
axes[1].bar(x, data_637, color=colors)
axes[1].set_title("$m/z$ = 637.176", size = 15)
axes[1].set_ylabel("Peak height" , size = 15)
axes[1].set_xticks(x)
axes[1].set_xticklabels(labels)


plt.tight_layout()

# SVG保存
plt.savefig("D:/発表関連/原稿/peak_height_comparison.svg", format="svg")

plt.show()

# %% [code]
import matplotlib.pyplot as plt
import numpy as np

# データ
labels = ["$G. uralensis$ root", "$G. uralensis$ leaf/stem", "$G. glabra$ root", "$G. glabra$ leaf/stem"]

data_505 = [10373, 32, 51386, 64]
data_637 = [10785, 17, 22403, 57]

x = np.arange(len(labels))

# カラー
colors = ["#FF5733", "#4DAA64", "#9E0000", "#005F3C"]


# -------- 505 --------
plt.figure(figsize=(8,6))
plt.bar(x, data_505, color=colors)
plt.title("$m/z$ = 505.134", size=15)
plt.ylabel("Peak height", size=15)
plt.xticks(x, labels, size=11)

plt.tight_layout()
plt.savefig("D:/発表関連/原稿/peak_height_505.svg", format="svg")
plt.show()

# -------- 637 --------
plt.figure(figsize=(8,6))
plt.bar(x, data_637, color=colors)
plt.title("$m/z$ = 637.176", size=15)
plt.ylabel("Peak height", size=15)
plt.xticks(x, labels, size=11)

plt.tight_layout()
plt.savefig("D:/発表関連/原稿/peak_height_637.svg", format="svg")
plt.show()
```
