## Supplementary Figures for "Stable isotope–assisted computational mass spectrometry reveals root-specific alkaloids in *Glycyrrhiza* species"

### Corresponding Author

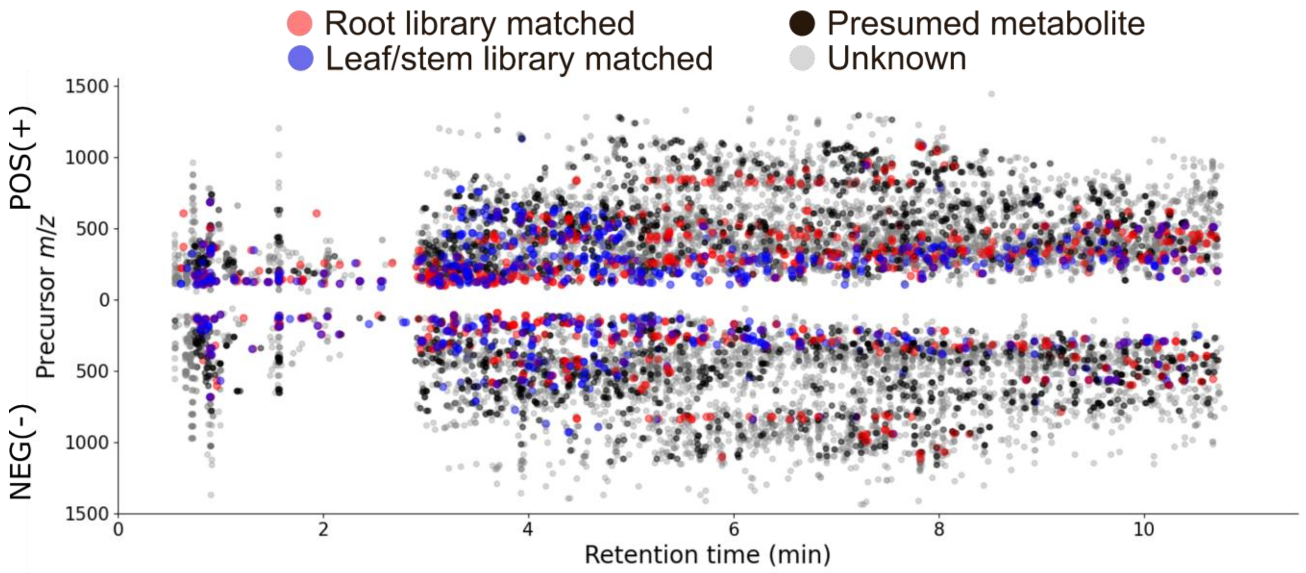


POS(+) | NEG(-)

| | *G.uralensis* root | *G.uralensis* leaf/stem | *G.glabra* root | *G.glabra* leaf/stem |
|---|---|---|---|---|
| MS/MS acquired | 2072 \| 1455 | 1555 \| 1210 | 2285 \|1621 | 1841 \| 1537 |
| Library matched | 361 \| 197 | 273 \| 129 | 353 \| 194 | 317 \| 194 |
| Presumed metabolite | 657 \| 540 | 330 \| 319 | 395 \|329 | 254 \| 236 |

**Supplementary Figure 1. Scatter plot of detected peaks.** Library-matched peaks were mapped in red (root) and blue (leaf/stem). Presumed ions are mapped in gray, and redundant ions are mapped in black.

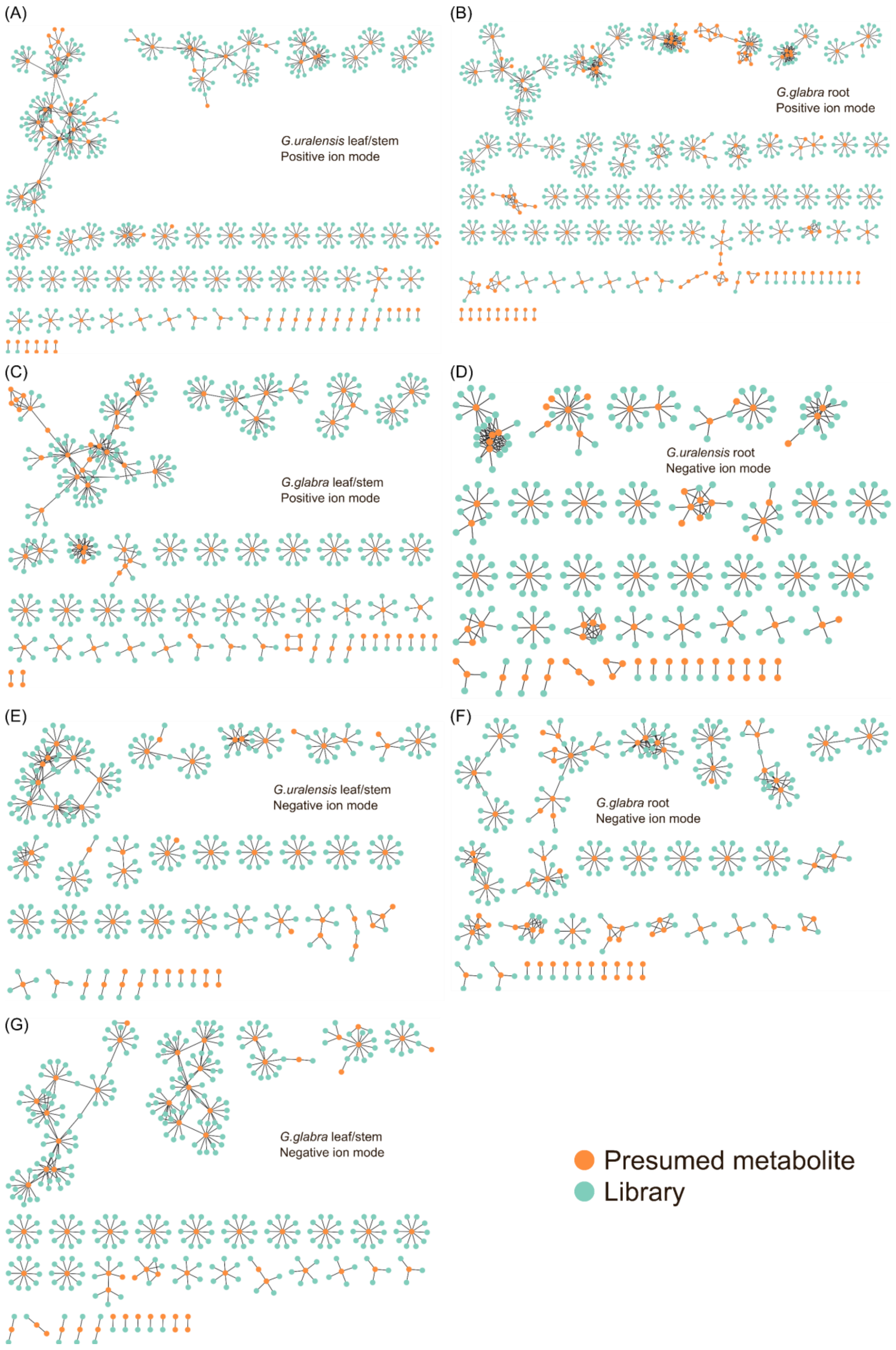


**Supplementary Figure 2. MS/MS molecular networks constructed using data clean-up ions and reference library spectra.** (A) *Glycyrrhiza uralensis* root, (B) *G. uralensis* leaf/stem, (C) *G. glabra* root, and (D) *G. glabra* leaf/stem in the positive ion mode. (E) *G. uralensis* root, (F) *G. uralensis* leaf/stem, (G) *G. glabra* root, and (H) *G. glabra* leaf/stem in the negative ion mode.

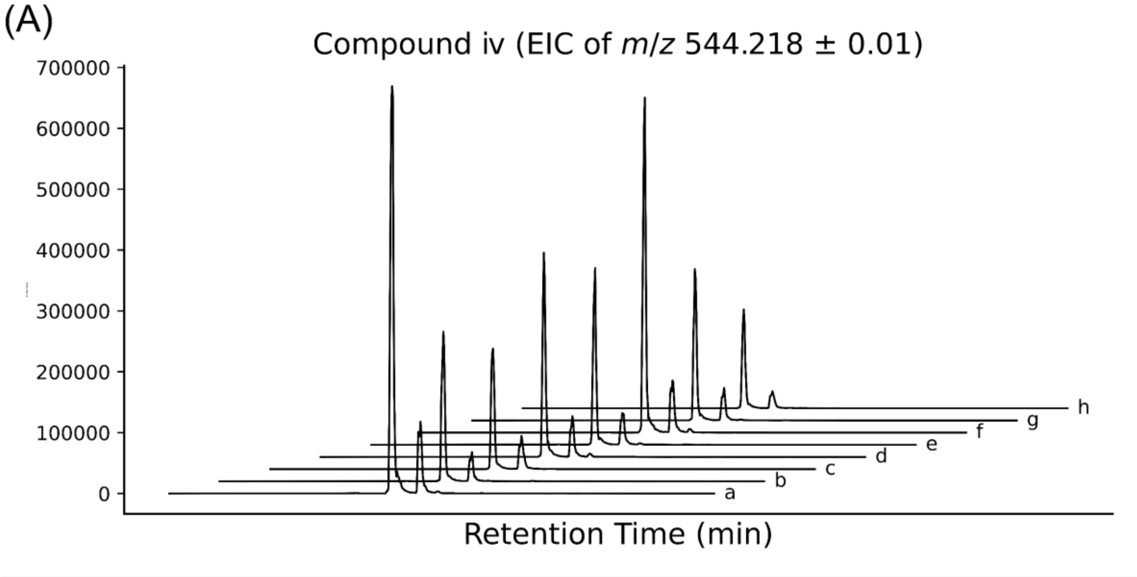


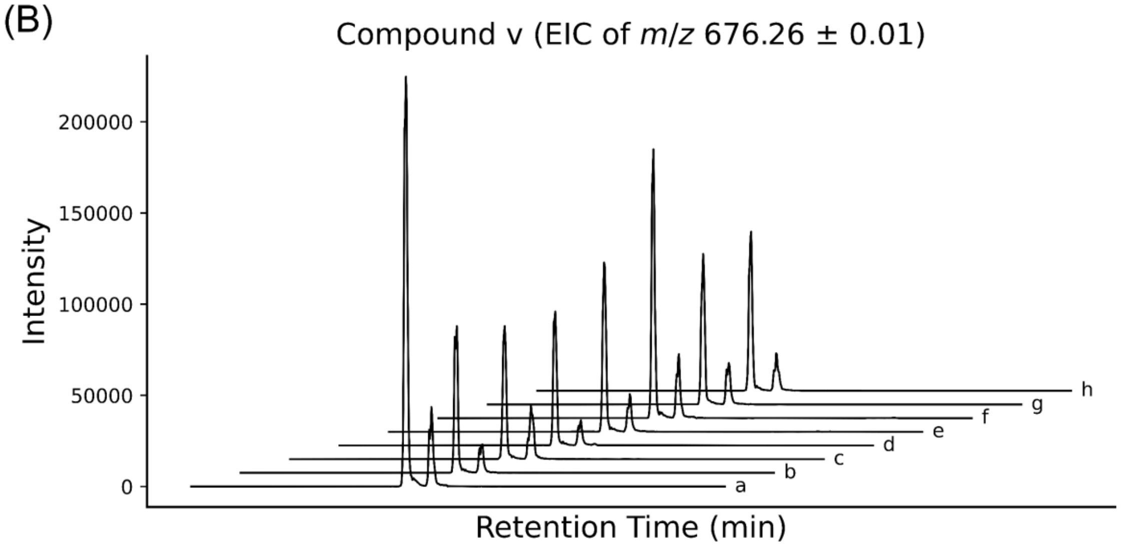


**Supplementary Figure 3. Extracted ion chromatograms (EICs) of compound (iv) (A) and compound (v). (B) obtained from different commercial licorice products**. (a) Tohoku (Uchida), (b) Tohoku-joh (Uchida), (c) Seihoku (Uchida), (d) Seihoku-joh (Uchida), (e) Seihoku-kizami (Tochimoto), (f) licorice (Tochimoto), (g) Licorice powder (Tochimoto), and (h) Licorice powder (peeled) (Tochimoto).

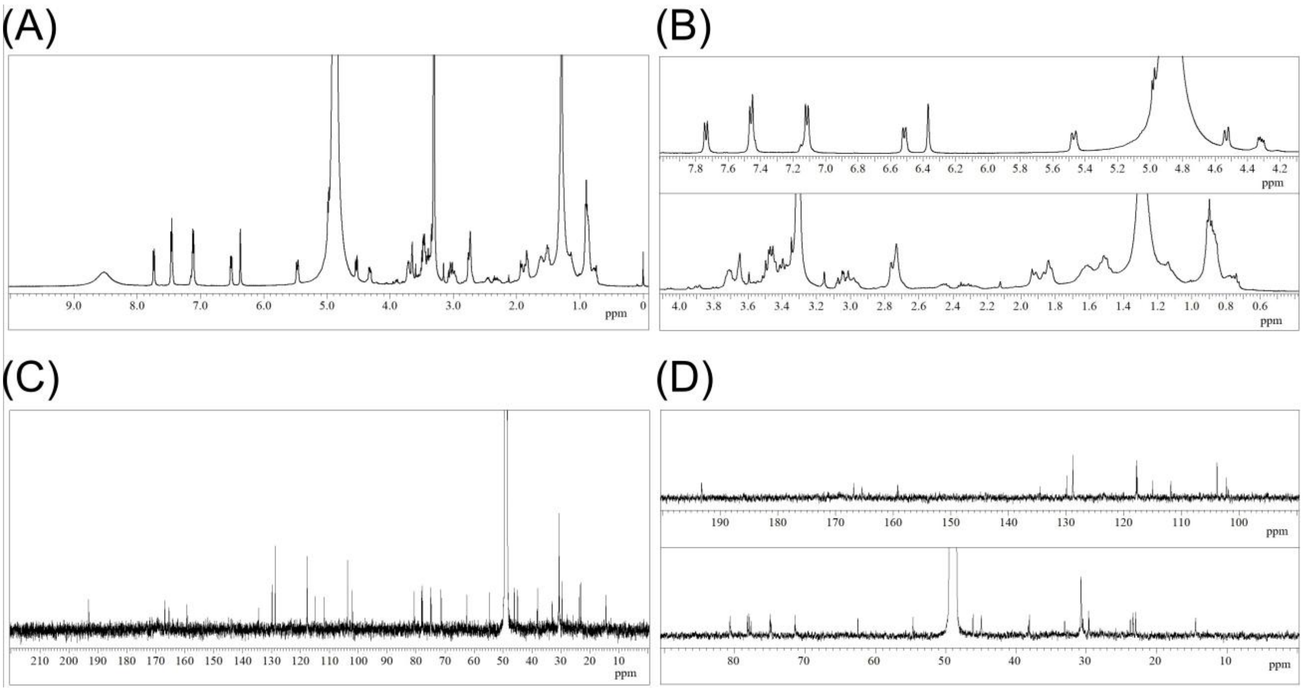


**Supplementary Figure 4. NMR spectrum data for compound (iv).** (A) $^{1}$H NMR spectrum. (B) Expanded view of the spectrum. Compound (iv): $^{1}$H NMR (500 MHz, $CD_3OD$, $\delta_H$ in ppm, TMS as internal standard) $\delta_H$ 5.47 (*d*, J = 12.5 Hz, H-2), 3.04* (*m*, H-3a), 2.75* (*m*, H-3b), 7.74 (*d*, *J* = 8.6 Hz, H-5), 6.51 (*d*, J = 8.6 Hz, H-6), 6.37 (*s*, H-8), 7.46 (*d*, *J* = 8.2 Hz, H-2′, H-6′), 7.11 (*d*, *J* = 8.2 Hz, H-3′, H-5′), 4.98 (*d*, *J* = 7.3 Hz, H-1″), 3.47* (*m*, H-2″), 3.49* (*m*, H-3″), 3.40 (*m*, H-4″), 3.71 (*m*, H-5″), 4.53 (*d*, *J* = 11.5 Hz, H-6″a), 4.32 (*d*, *J* = 11.5 Hz, H-6″b), 3.46* (*m*, H-11), 1.92 (*m*, H-12a), 1.52 (*m*, H-12b), 1.90* (*m*, H-13a, H-14a), 1.60* (*m*, H-13b, H-14b), 3.36* (*m*, H-15a), 2.98* (*m*, H-15b), 2.72* (*m*, H-17). Compound (v): $\delta_H$ 5.05 (*d*, *J* = 7.0 Hz, H-1‴), 3.65* (*m*, H-2‴), 4.05 (*dd*, *J* = 9.5, 6.3 Hz, H-4‴a), 3.80 (*dd*, *J* = 9.5, 6.3 Hz, H-4‴b), 3.53* (*m*, H-5‴). (C) $^{13}$C NMR spectrum. (D) Expanded view of the spectrum. Compound (iv): $^{13}$C NMR (125 MHz, $CD_3OD$, $\delta_C$ in ppm, referenced to $CD_3OD$ at 49.0 ppm) $\delta_C$ 80.7 (C-2), 45.0 (C-3), 193.2 (C-4), 129.9 (C-5), 111.8 (C-6), 166.9 (C-7), 103.8 (C-8), 165.4 (C-9), 115.0 (C-1′), 134.5 (C-2′), 128.8 (C-3′, C-5′), 117.8 (C-4′, C-6′), 159.3 (C-4′), 101.9 (C-1″), 74.9 (C-2″), 77.7 (C-3″), 71.4 (C-4″), 75.0 (C-5″), 65.3 (C-6″), 54.7 (C-11), 30.5* (C-12), 23.3* (C-13, C-14), 46.1 (C-15), 38.2 (C-17), 172.0 (C-18). Compound (v): $\delta_C$ 102.2 (C-1‴), 78.2 (C-2‴), 80.6* (C-3‴), 75.5 (C-4‴), 66.0 (C-5‴). *Overlapped peaks, identified from COSY, HSQC and HMBC.

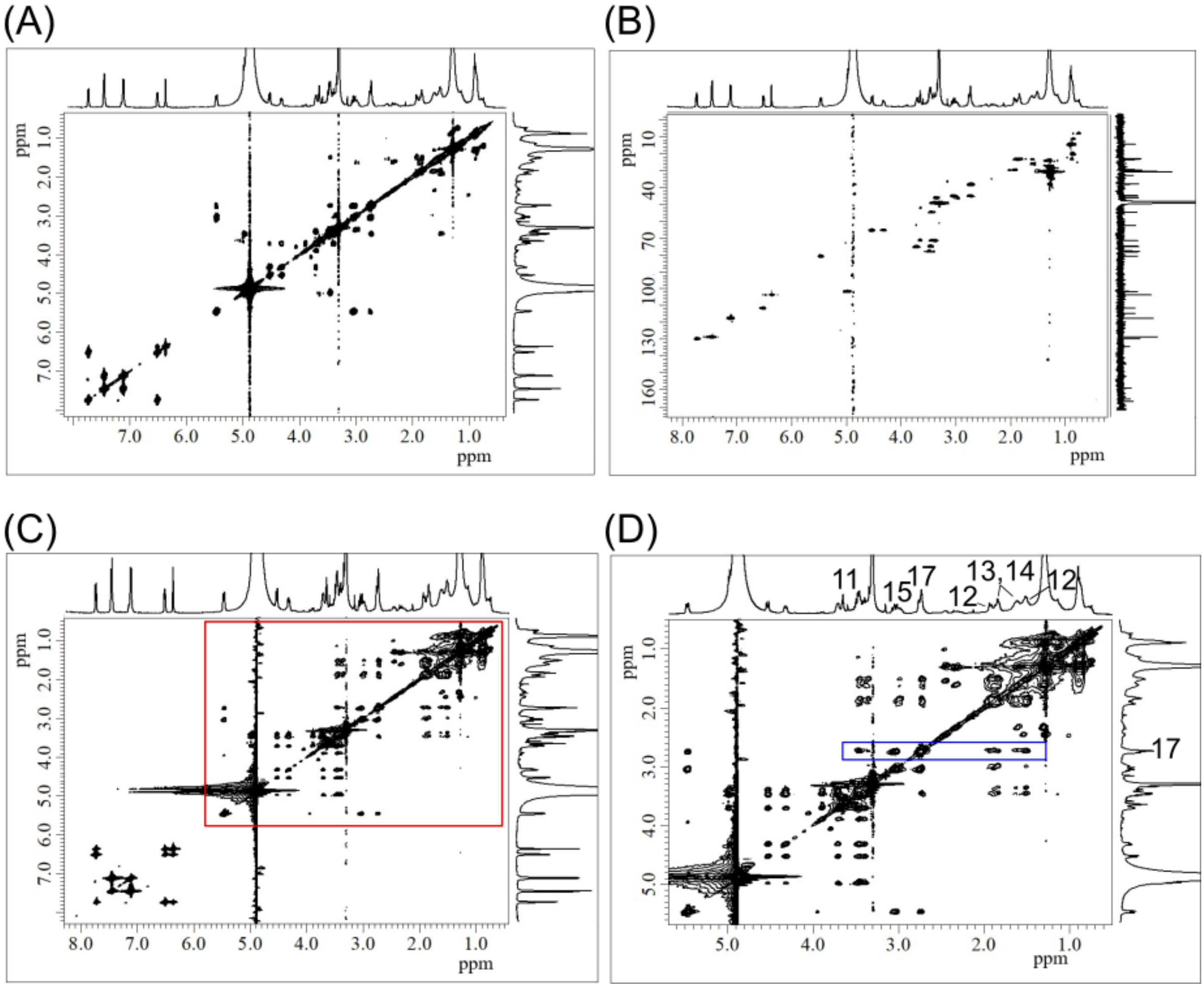


**Supplementary Figure 5. COSY, HSQC, and TOCSY NMR spectra for compound (iv).** (A) COSY (Correlation Spectroscopy), (B) HSQC (Heteronuclear Single Quantum Coherence), and (C) TOCSY (Total Correlation Spectroscopy) spectra. (D) Expanded view of the region highlighted by the red box. The correlations highlighted by the blue box suggest a continuous TOCSY spin system extending from H-17 through H-11, H-12, H-13, and H-14 to H-15, consistent with the presence of a six-membered ring structure (iv).

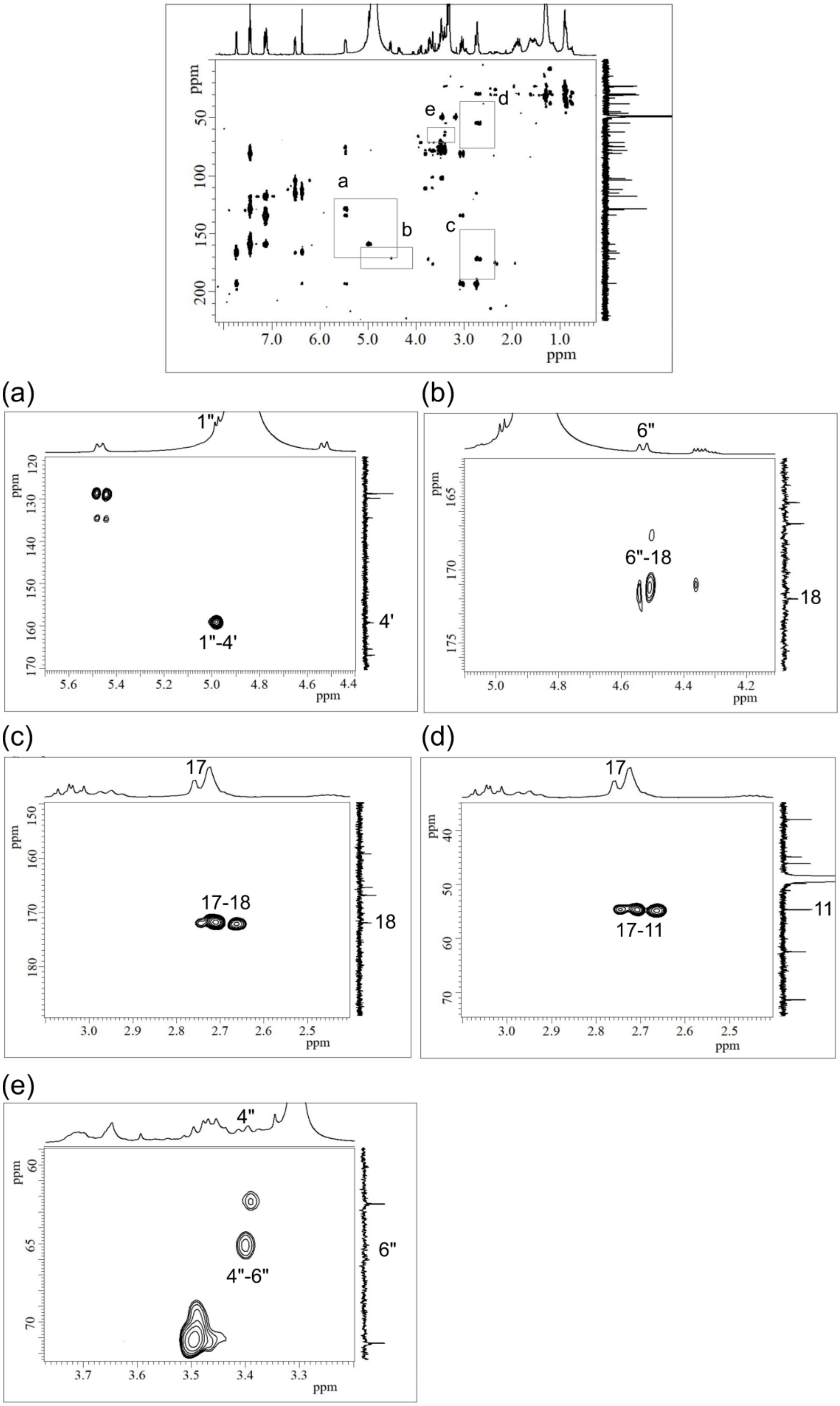


**Supplementary Figure 6. HMBC (Heteronuclear Multiple Bond Correlation) spectrum.** Regions showing key correlations for structural elucidation are highlighted by boxes and shown as expanded views: (a) 1″-4′, (b) 6″-18, (c) 17-18, (d) 17-11, and (e) 4″-6″.

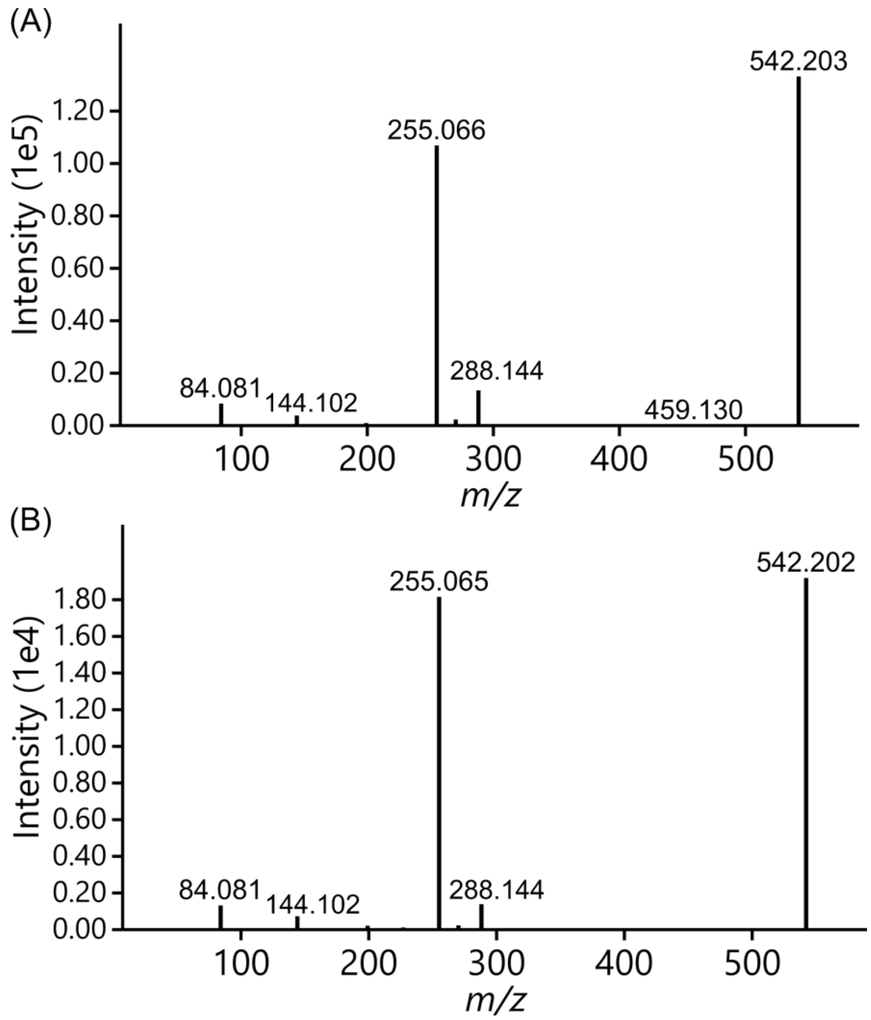


**Supplementary Figure 7. MS/MS spectra of isoflavone base; 2O; O-Hex; HA, generated by mixing 6”-O-malonyldaidzin with 1-piperidine at (A) 37 °C and (B) room temperature.**

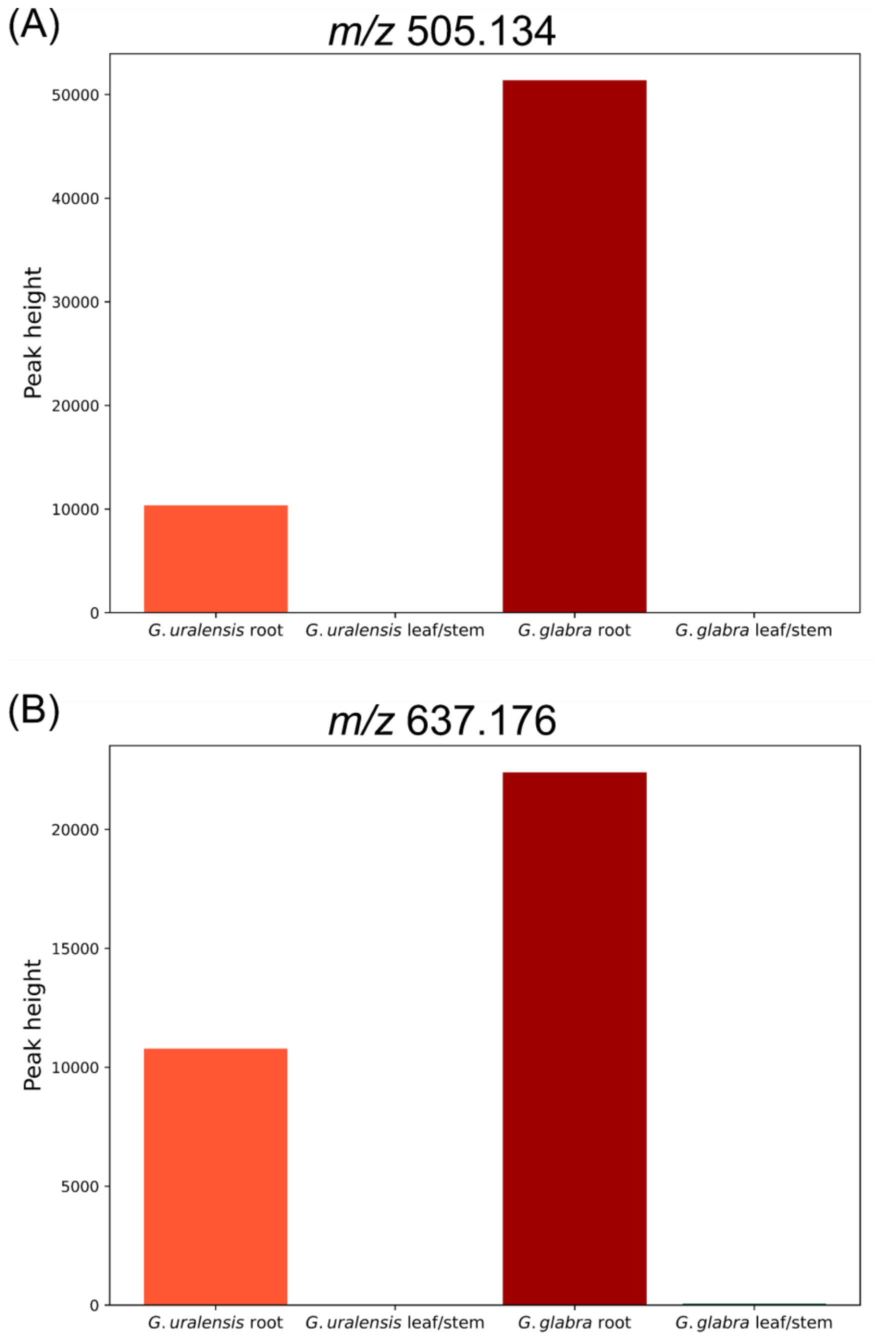


**Supplementary Figure 8. Distribution of (A) flavanone base; 2O; O-MHex and (B) flavanone base; 2O; O-Pen; O-MHex.**

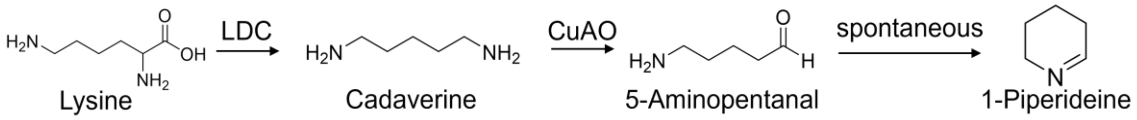


**Supplementary Figure 9. The proposed biosynthetic pathway for 1-piperideine from lysine is mediated by lysine decarboxylase (LDC) and copper amine oxidase (CuAO).**
